# New AAV9 engineered variants with enhanced neurotropism and reduced liver off-targeting in mice and marmosets

**DOI:** 10.1101/2023.06.27.546696

**Authors:** Serena Gea Giannelli, Mirko Luoni, Benedetta Bellinazzi, Angelo Iannielli, Jinte Middeldorp, Ingrid Philippens, Jakob Körbelin, Vania Broccoli

**Author notes:** **Lead contact: Vania Broccoli** - Stem Cells and Neurogenesis Unit, Division of Neuroscience, San Raffaele Scientific Institute, Via Olgettina 58, 20132 Milan, Italy. These authors share first authorship (Serena Gea Giannelli, Mirko Luoni).

## Abstract

Adeno-Associated Virus 9 (AAV9) is a delivery platform highly exploited to develop gene-based treatments for neurological disorders given its low pathogenicity and brain tissue tropism. However, the efficacy of this vector is dampened by its relatively low efficiency to cross the adult blood-brain barrier (BBB) and inherent targeting to the liver upon intravenous delivery. We generated a new peptide display library starting from a galactose binding-deficient AAV9 capsid and selected two new AAV9 engineered capsids, named AAV-Se1 and AAV-Se2, with an enhanced targeting in mouse and marmoset brains after intravenous delivery. Interestingly, the loss of the galactose binding strongly reduced the undesired targeting to peripheral organs, and above all liver, while not compromising the transduction of the brain vasculature. However, we had to reconstitute the galactose binding in order to efficiently infect non-endothelial brain cells. Thus, the combinatorial actions of the galactose-binding domain and the installed exogenous displayed peptide are crucial to enhance BBB crossing together with brain cell transduction. We also identified Ly6C1 as primary receptor for AAV-Se2 which is a Ly6A homologue highly expressed in the brain endothelial cells. This study describes a new strategy to select neurotropic AAV9 variants and identifies two novel capsids with high brain endothelial infectivity and extremely low liver targeting based on manipulating the AAV9 galactose binding domain.

## Introduction

Adeno-associated virus (AAV) vectors are the most commonly employed delivery systems for *in vivo* gene therapy in preclinical and clinical studies^1–3^. AAV vectors are composed of a protein capsid which shields a single-stranded DNA genome and determines the vector tropism. Natural serotypes offer a suite of delivery platforms with multiple opportunities for tissue and organ targeting^4^. However, all of them have a significant tropism for the liver when administrated intravenously limiting the transduction efficiency in other organs and raising important concerns of toxicity^5–7^. The definition of the AAV capsid structure at high-resolution, and identification of their cellular receptors and intracellular pathways have provided the necessary knowledge for introducing targeted modifications aiming to re-configurate the viral behavior *in vivo* as for instance its tissue tropism, stability or immunity^8, 9^. To this end, several AAV library-based approaches have been elaborated to isolate new variants with the desired property through an iterative selection *in vitro* or *in vivo* known as directed evolution. This strategy has proven very powerful yielding engineered capsids with compelling new or improved features. Diverse modalities have been exploited for rational capsid engineering, as in particular, single amino acid substitutions, capsid chimeras through DNA shuffling, or targeted insertion of random peptides in key positions controlling capsid-cell receptor interactions, an approach referred to as random peptide display^10–12^. This last strategy was pioneered by two German groups who implanted a random 7-amino acid sequences in positions R588 and N587, respectively, of the AAV2-VP1 in order to introduce high sequence diversity on the superficial domain of the capsid to maximize the engagement with new cell receptors^13, 14^. Importantly, this domain is mainly responsible for the binding of AAV2 to its natural receptor heparan sulphate^15^. In fact, the integration of the peptide reduced by at least 70% the binding to heparin respect to the unmodified AAV2^13^. Thus, this strategy, from one side significantly reduced the AAV2 natural binding while, and on the other side, surveyed new interactions through the display of a random sequence. This design coupled with multiple rounds of capsid selection *in vivo*, enabled for the isolation of variants with superior tropism respect to the parental AAV2 for coronary artery, pulmonary and brain endothelial cells^13, 16, 17^, lung^18^, liver^19^ and cardiomyocytes^20^ among others. With the same rational, a random heptapeptide sequence was inserted in the AAV9 capsid in position A589 corresponding to the analogue domain of the AAV2 capsid^21, 22^. However, differently from AAV2, this domain is not responsible for AAV9 binding to galactose, its natural primary cell receptor^2–25^. Thus, this peptide display system diversifies an important domain on the surface of the AAV9 capsid, but its interactions should complement or counteract the binding specificity dictated by the galactose engagement which remains in place yet. Using an analogous library equipped with a Cre-dependent selectable cassette, the PHP.B viral family was isolated with a superior ability to cross the BBB and transduce the central and peripheral nervous systems^26, 27^. However, this behavior is not conserved in other mammals since their transduction was found to depend by the binding to the rodent-specific Ly6A endothelial receptor^28–31^. Remarkably, in vivo selection of AAV9 peptide display libraries led also to the identification of RGD motif-containing capsids with a superior efficiency and selectivity for muscle tissues after intravenous injections in mice and non-human primates (NHPs)^32, 33^. Thus, AAV engineered libraries coupled with effective selection strategies have provided an invaluable system to isolate new capsids with selected traits tailored for specific applications. Herein, we generated a new display library using the AAV9 galactose binding-mutant capsid to abrogate its natural tropism and redirect it to alternative cell receptors. Iterative *in vivo* selection for superior neurotropic viruses after intravenous delivery, identified two novel variants which outperformed the parental natural capsid and maintained high transduction efficiency in both C57BL/6 and BALBc mice. Moreover, both engineered capsids showed superior gene transfer capability in adult marmosets and human neuronal cultures with respect to the unmodified AAV9. Interestingly, we showed that the lack of galactose-binding suppressed the off-target transduction in the liver, without compromising the targeting to the brain vasculature.

## Results

### Generation and screening of a new AAV9 peptide display library

In order to maximize the targeting diversity of the AAV9, we reasoned that removing the galactose binding of the capsid would facilitate the identification of new interactions between the random display peptides and their cellular targets. The AAV9 capsid contains a small galactose binding pocket and point mutations in this domain reduce its binding compromising its transduction efficiency in tissues^23–25^. We introduced the W503A mutation which was described to virtually eliminate galactose binding preventing the viral transduction in cells^25^. Then, we inserted a randomized sequence of 7 amino acids (7-mer) between amino acids 588 and 589 of the mutant W503A VP1 variant (Figure 1A). Based on the cloning strategy two additional amino acids, a glycine and alanine, were flanking the random sequence. We speculated that given their small hindrance and neutral charge, these amino acids would not represent a significant impediment to the binding, but rather enhance the flexibility of the loop created by the random sequence. The diversity of the produced plasmid library was 1 × 10^8^ clones per library as assessed by targeted deep-sequencing. Next, we transferred the plasmid library into AAV producing cells with a minimal amount of plasmid per cell to minimize that multiple plasmids would be incorporated into single producer cells with the high risk to produce chimeric capsids. Sequencing of infected clones indicated that the diversity of the viral library was comparable to that of the plasmid library ensuring an efficient and unbiased production of the AAV library.

**Figure 1:**
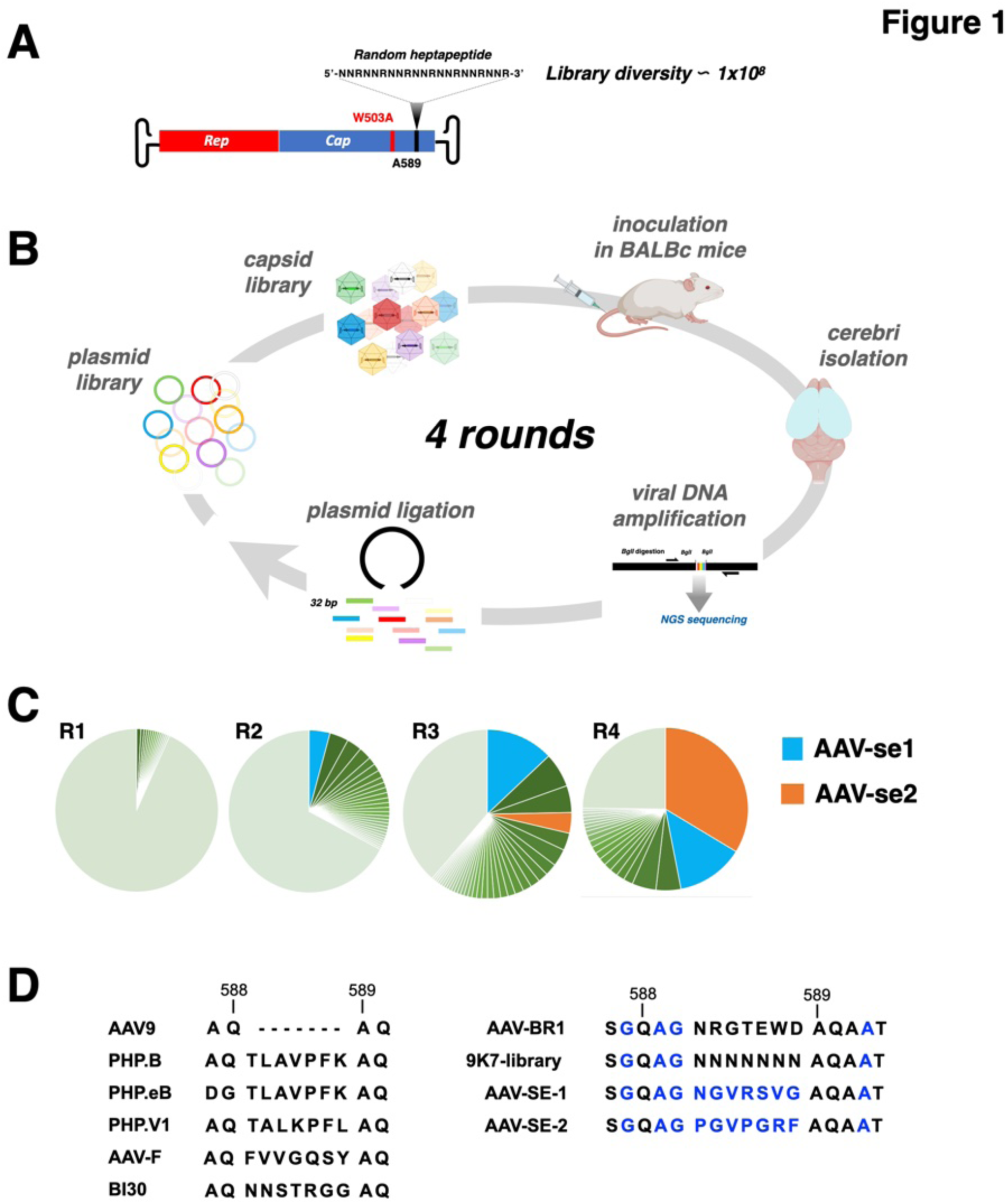
Outline of AAV9 peptide display library selection procedure and NGS sequencing results. **A)** Illustration of AAV9 construct indicating the distinctive features of our library: the W503A mutation in the *Cap* cistron that amperes galactose binding and the position A589 where the heptapeptide display library was inserted. The heptapeptide library was encoded by a 21nt, whose codons in the third position presents only with a G or an A (R). **B)** Illustration of the main procedures undertaken during each round of library screening. **C)** Pie charts indicate the frequency of each different 7mer sequences determined by NGS. In the brain, the highest ranking 7mer at the 3^rd^ round (R3) was named AAV-Se1 (Blue), which upon a 4^th^ round (R4) was overcome by another variant, named AAV-Se2 (Orange). **D)** AAV-Se1 and -Se2 heptamers and contiguous residues are compared to the parental library sequence, native AAV9, as well as various other AAV9- (PHP.B, PHP.eB, PHP.V1, AAV-F and CAP-B130) and AAV2- (AAV-BR1) based capsid variants.

We, next, sought to exploit this library to identify new AAV9 variants acquiring the ability to cross the BBB in adult BALBc mice. This mouse strain has proven to be resistant to brain transduction by the PHP.B family of capsids since their putative receptor Ly6A is mutated and not exposed to the cell membrane^28–31^. Thus, these mice offered a convenient and straightforward system to prevent the selection of Ly6A-binding capsids to begin with. A single dose of 2 × 10^11^ vector genomes (vg) of the display peptide library was injected into the tail vein of BALBc adult mice (Figure 1B). 3 weeks after viral inoculation, the brain was isolated, cerebral cortices dissected and lysated to harvest the DNA of the viral particles that successfully homed into the brain (Figure 1B). The relevant part of the capsid gene was amplified by PCR, re-cloned into a new library, and the selection process was then repeated for three additional rounds (Figure 1B). After each round, deep-sequencing of the viral DNA amplicon was performed to trace the progressive enrichment of viral sequences with preferential targeting to the brain tissue (Figure 1B). After the first two rounds of selection, 5 out of the 7 residues had strongly enriched one specific amino acid (Figure S1). After the third round the most enriched peptide was NGVRSVG (13% of total reads), however a new, and clearly distinct, motif became apparent to the following and last cycle, with the sequence PGVPGRF (34% of total reads) (Figures 1C,D). We named the two new variants as AAV-A503-Se1 and AAV-A503-Se2 in the order they appeared along the different rounds of selection. The presence of the W503A mutation did not alter significantly the yield of viral production of either the library or the selected variants.

### Brain transduction pattern of AAV-A503-Se1 and AAV-A503-Se2 in C57BL/6 and BALBc mice

We generated a reporter vector containing the ZsGreen, which ensures a brighter fluorescence compared to GFP, under the control of the CBA promoter and packaged it into the AAV-A503-Se1 and AAV-A503-Se2 capsids (Figure 2A). In addition, we generated an AAV9-A503 mutant capsid as a pertinent negative control. In fact, C57BL/6 mice injected with the AAV9 capsid mutant at the dose of 1 × 10^11^ vg/mouse failed to show ZsGreen staining in the brain confirming a loss of transduction as expected (Figure 2B). Conversely, both AAV-A503-Se1/2 capsid variants at the same dose sustained a diffuse transgene expression throughout the brain (Figure 2B). Importantly, ZsGreen expression was comparable in both C57BL/6 and BALBc strains of mice for both capsid variants suggesting that their transduction was independent from the PHP.B receptor Ly6A (Figure 2B). In particular, we noted that mice treated with either capsid variant showed intense ZsGreen fluorescence in the vasculature throughout the cortices (Figure 2B). Double immunofluorescence staining between the ZsGreen and the endothelial-specific marker CD31 (Pecam1) showed over 80% of productive viral transduction of endothelial cells throughout the brain for both AAV-Se variants. In contrast, only a minority of CD31^-^ cells, all together below 8%, in the brain parenchyma expressed the viral transgene with either variant in any of the two mouse strains (Figure 2C-E). In line with previous results, viral genome quantifications by qPCR revealed a 12- and 16-fold increase from lysates of brains transduced, respectively, with AAV-A503-Se1 and -Se2 respect to the AAV9-A503 (Figure 2F). These findings indicate that the AAV-A503-Se variants acquired the ability to efficiently target the brain by transducing mostly its vasculature while being highly inefficient to infect other brain cells.

**Figure 2:**
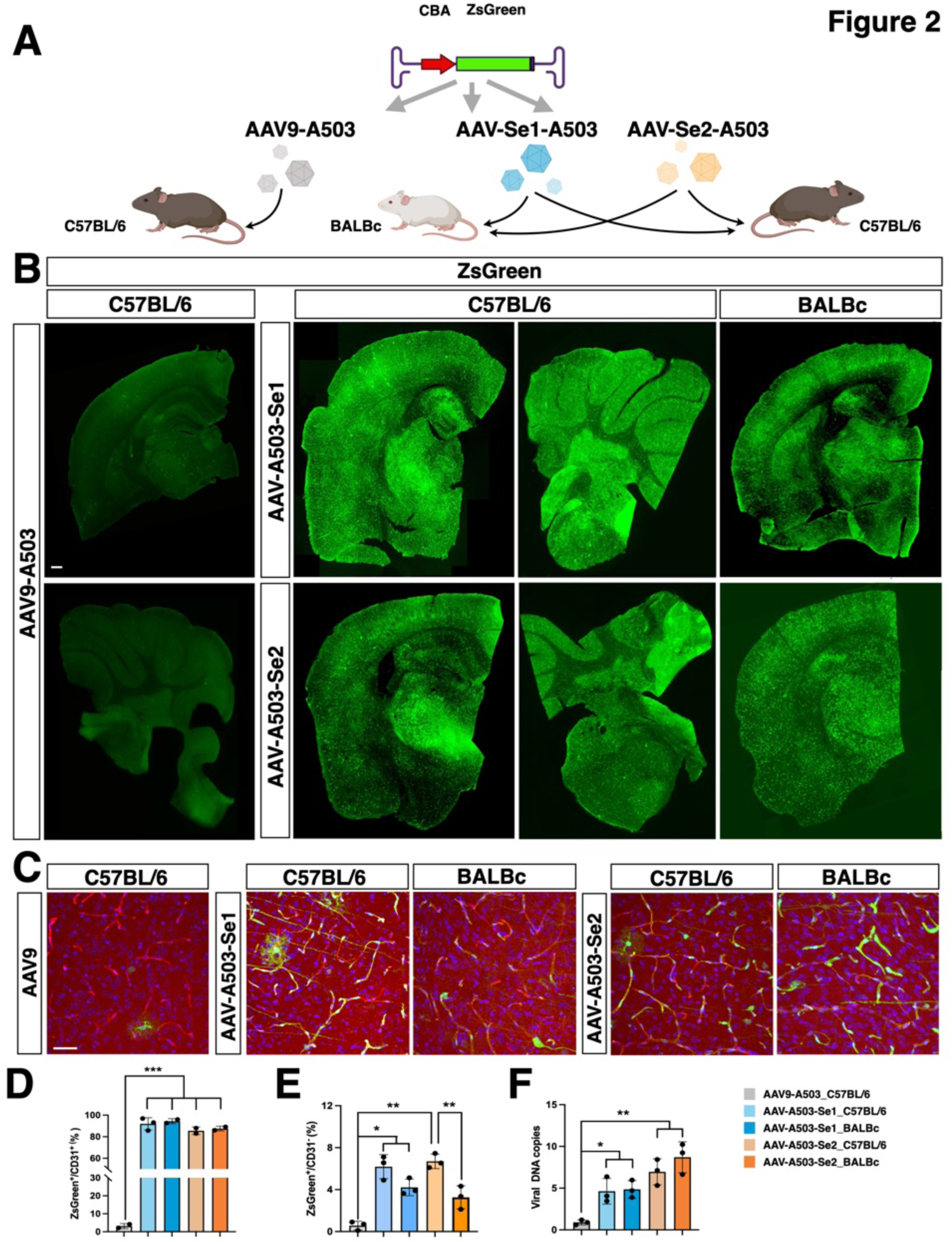
AAV-A503-Se1 and AAV-A503-Se2 exhibit a high brain tropism selectively for endothelial cells in both C57BL/6 and BALBc mice. **A)** Illustration of the experimental setting. Wild-type C57BL/6 and BALBc mice are systemically injected with AAV9-A503, AAV-A503-Se1 and AAV-A503-Se2 carrying the ZsGreen transgene under the control of the CBA strong constitutive promoter (experimental viral dose: 1 × 10^11^ vg/mouse, n = 3 animals for each group). **B)** Low magnification immunostaining for ZsGreen, in cerebrum and cerebellum derived from animals treated with AAV9-A503, AAV-A503-Se1 and AAV-A503-Se2. Scale bar, 200 µm. **C)** High magnification immunostaining for ZsGreen and CD31 (in red) in cortex derived from animals treated with AAV9-A503, AAV-A503-Se1 and AAV-A503-Se2. Scale bar, 100 µm. **D)** Quantitative assessment of virally transduced (ZsGreen^+^) endothelial cells (CD31^+^) in brain derived from AAV9-A503, AAV-A503-Se1 and AAV-A503-Se2 treated C57BL/6 and BALBc mice (n = 2-3 animals per group). **E)** Percentage of non-endothelial cells (CD31^-^) transduced by the AAV-A503-Se1 and AAV-A503-Se2 in C57BL/6 and BALBc mice (n = 3 animals per group). **F)** Viral DNA vector quantification by qRT-PCR in cortexes derived from AAV9-A503, AAV-A503-Se1 and AAV-A503-Se2 treated C57BL/6 and BALBc mice. The results are reported as the fold change of viral ZsGreen DNA in mice treated with AAV-A503-Se1 and AAV-A503-Se2 relative to mice treated with AAV9-A503 (n = 3 animals per group). Values are mean ± SD. *p < 0.05; **p < 0.01, ***p < 0.001. Statistical analysis is performed using one-way ANOVA followed by Tukey post-test.

### High and diffuse brain transduction of AAV-Se capsid variants by restoring the galactose binding capability

We reasoned that a possible explanation for the lack of the AAV-A503-Se capsids to transduce brain cells beyond the vasculature was their inability to bind galactose. To assess if this was the case, we reverted the Alanine into Tryptophan in position 503 of VP1 in both AAV-Se capsids to restore their capacity to bind to galactose (Figure 3A). Then, we packaged a single-stranded ZsGreen-expressing vector in either capsid, and delivered 1 × 10^11^ vg of viruses by intravenous injections in adult mice. As reference controls, we produced AAV9 and AAV-PHP.eB viruses packaging the equivalent reporter and inoculated in mice as executed for the AAV-Se capsids. Three weeks post-transduction, ZsGreen distribution was assessed within the brains in both C57BL/6 and BALBc mice. Interestingly, wild-type AAV-Se1/2 showed a significantly more diffuse tropism in the brain respect to their mutant analogues and the unmodified AAV9 (Figure 3B). Although the general level of distribution of the AAV-Se1/2 viruses appeared comparable to that of PHP.eB in C57BL/6 mice, their pattern of infection was not significantly changed in BALBc mice as it occurred with the PHP.eB (Figure 3B). qPCR quantifications of viral genomes in brain lysates confirmed a substantial comparable transduction efficiency of AAV-Se1/2 in either mouse strain, which resulted highly superior to that of standard AAV9 and not significantly different from the gene transfer levels reached by the AAV-PHP.eB (Figure 3C). To determine the identity of the brain cells transduced by AAV-Se1/2, we performed immunohistochemistry for specific markers of neurons (NeuN), astrocytes (Sox9) and endothelial cells (CD31), three weeks after intravenous transduction of adult C57BL/6 and BALBc mice (Figure 4). Wild-type AAV-Se1 maintained a tropism mainly selective for CD31^+^ brain endothelial cells in both mouse strains (92% ± 3% in C57BL/6 and 86% ± 3% in BALBc). Non-endothelial cells infected by AAV-Se1 were very rare and predominantly astrocytes in the brain parenchyma (0.4% ± 0.1% and 0.2% ± 0.05% of NeuN^+^ neurons in C57BL/6 and BALBc, respectively; 0.7% ± 0.2% and 0.6% ± 0.3% of Sox9^+^ astrocytes in C57BL/6 and BALBc, respectively) (Figure 4). In contrast, wild-type AAV-Se2-mediated transgene expression was found more broadly in the brain parenchyma. In fact, AAV-Se2 gene transfer in brain cells was significantly more efficient with respect to the parental AAV9. AAV-Se2 sustained an increased transduction of neuronal and glial cells compared to the AAV9 by a factor of 7 and 5, respectively (Figure 4). Of note, AAV-Se2 targeted more astrocytes (42.38% ± 5.73%) than PHP.eB (34.21% ± 4.25%), and the reverse was true for neurons (19.67% ± 4.2% for AAV-Se2 and 38.29% ± 3.16% for PHP.eB), suggestive of a slightly different tropism between those two capsids in C57BL/6 mice (Figure 4). No appreciable region-to-region variation was observed in AAV-Se1/2-mediated gene transfer in terms of cell type specificity and total infection rate suggesting a consistent and homogeneous transduction profile across the entire brain. Finally, we compared the infectivity rates of the parental AAV9 with its capsid variants on primary mouse neuronal cultures in the dish (Figure S2). Interestingly, wild-type AAV-Se1/Se2 showed a significantly higher transduction efficiency on 10 days *in vitro* mouse primary neuronal cultures compared to both PHP.eB and unmodified AAV9 showed (AAV9 39% ± 1.8%; PHP.eB, 38% ± 3%; AAV-Se1, 88% ± 3%; AAV-Se2 81% ± 6% of Map2^+^ neurons) (Figure S2). Conversely, the A503 mutant capsids were incompetent to gene transfer with the exception of the AAV-A503-Se1 which sustained a diffuse infection although with reduced levels of ZsGreen expression (AAV-A502-Se1, 42% ± 1.5% of Map2^+^ neurons) (Figure S2).

**Figure 3:**
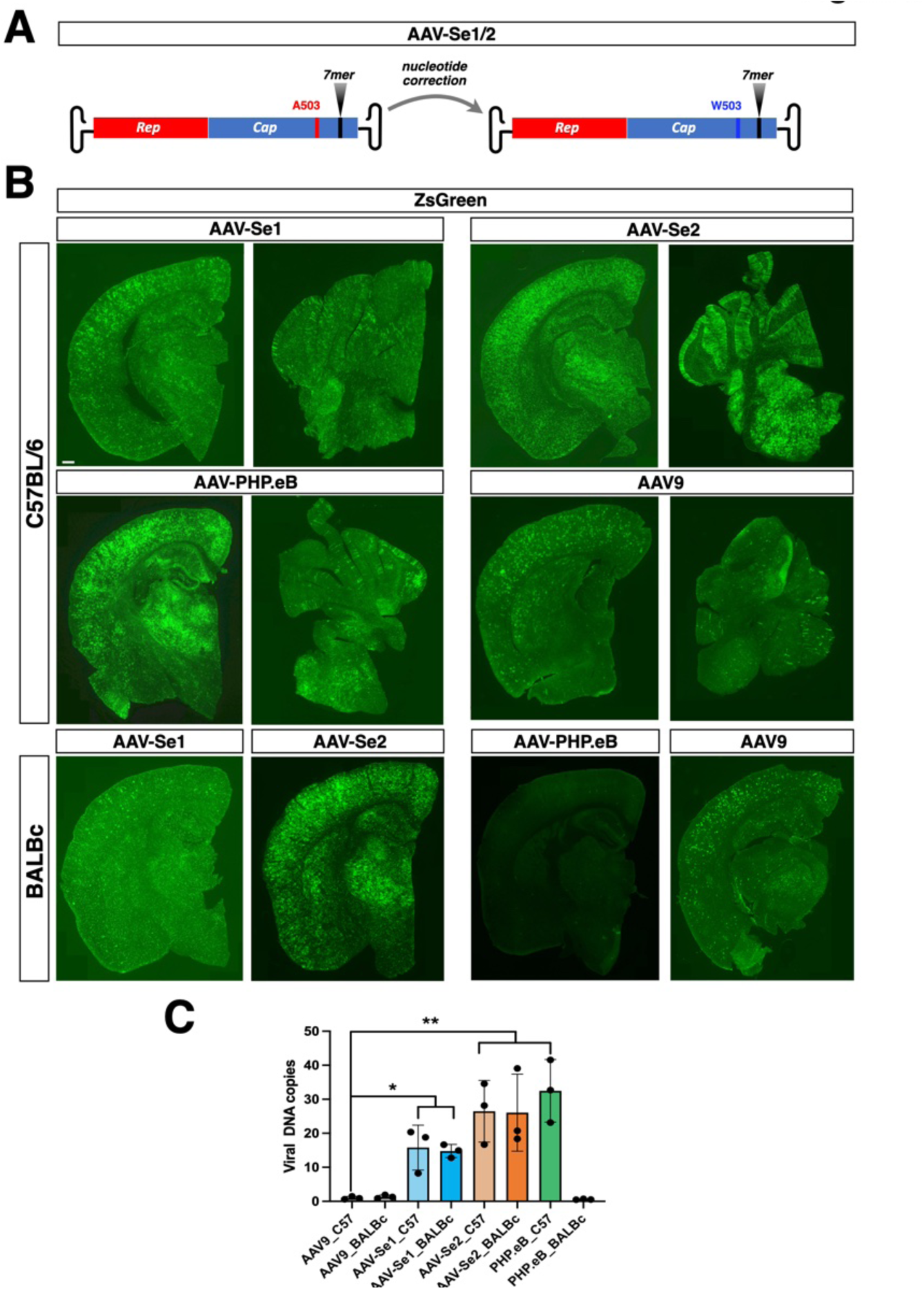
Restoring galactose binding reinforces the AAV-Se1/2 infectivity in both C57BL/6 and BALBc mice. **A)** Graphical illustration of the restoration of the galactose binding (A503W amino acid change). **B)** Low magnification immunostaining for ZsGreen in cerebrum and cerebellum derived from animals treated with AAV9, AAV-PHP.eB, AAV-Se1 and AAV-Se2. Scale bars, 200 µm. **C)** Viral DNA vector quantification by qRT-PCR in cortices derived from AAV9, AAV-PHP.eB, AAV-Se1 and AAV-Se2 treated C57BL/6 and BALB/c mice. The results are reported as fold change of viral ZsGreen DNA in mice treated with AAV-PHP.eB, AAV-Se1, AAV-Se2 relative to mice treated with AAV9 (n = 3 animals per group). Values are mean ± SD. *p < 0.05, **p < 0.01. Statistical analysis is performed using one-way ANOVA followed by Tukey post-test.

**Figure 4:**
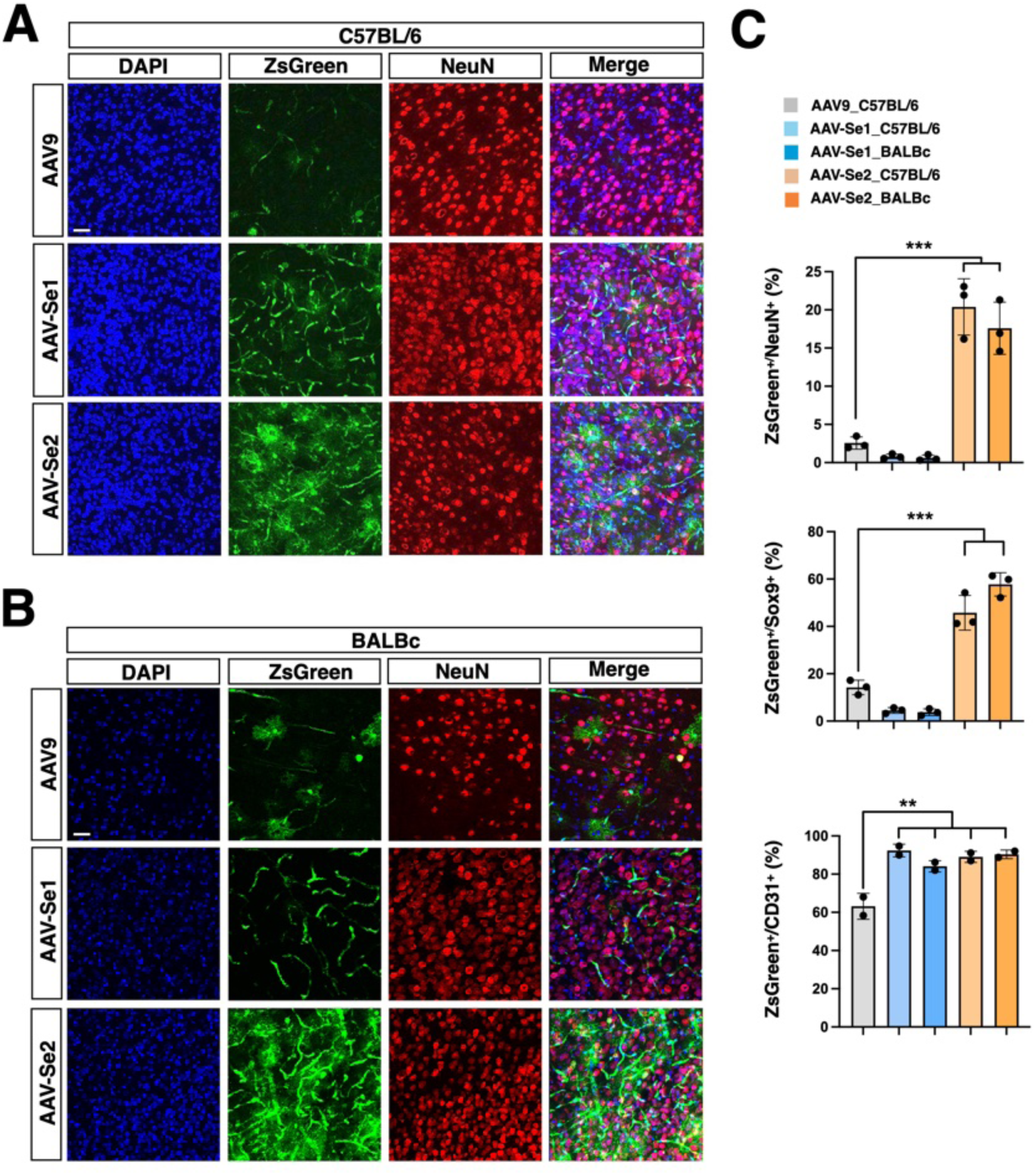
Cell-type specific analysis of the AAV-Se1/2 transduction pattern. **A, B)** High magnification images of immunostainings for ZsGreen and NeuN (red) in cortices derived from C57BL/6 (**A**) and BALBc (**B**) mice treated with AAV9, AAV-Se1 and AAV-Se2. Scale bars, 100 µm. **C)** Quantitative assessment of transduced (ZsGreen^+^) neurons (NeuN^+^), astrocytes (Sox9^+^) and endothelial cells (CD31^+^) in brain derived from AAV9, AAV-Se1 and AAV-Se2 treated C57BL/6 and BALBc mice (n = 2-3 animals per group). Values are mean ± SD of n = 3 independent experiments. **p < 0.01, ***p < 0.001. Statistical analysis is performed using one-way ANOVA followed by Tukey post-test.

### Distribution of AAV-Se viruses in peripheral organs after systemic delivery

In the fourth round of our original directed-evolution screening, we isolated a number of peripheral organs (liver, lung, heart, spleen, kidney, muscle), beyond the brain, and performed deep-sequencing of the viral amplicons to profile the off-target transduction propensity of the selected capsid variants. Interestingly, while the AAV-Se1 showed an enrichment in most of these organs, the AAV-Se2 was much less represented (Figure 5A). Next, we isolated the livers from C57BL/6 mice transduced with the AAV-Se1/2 and parental AAV9 viruses with or without the A503 mutation and expressing the ZsGreen expressing cassette, at the dose of 1 × 10^11^ vg/mouse. ZsGreen expression was barely detectable in livers transduced with the A503-mutant AAV9 and AAV-Se1/2 capsids indicating a general minimal tropism of these mutant viruses for this organ (Figure 5B). Moreover, viral DNA copy quantifications revealed an even further reduced transduction capability of the AAV-A503-Se capsids, and in particular of AAV-A503-Se2, respect to the parental AAV9 for the liver (Figure 5C). Correction of the A503 mutation in the AAV9 capsid enables to regain high tropism for the liver after intravenous administration in adult mice (Figure 5B). However, even the wild-type AAV-Se1/2 capsids maintained a significant reduced transduction rate in the liver compared to the parental capsid with 3- and 8-fold reduction for the AAV-Se1 and AAV-Se2, respectively (Figures 5B,C).

**Figure 5:**
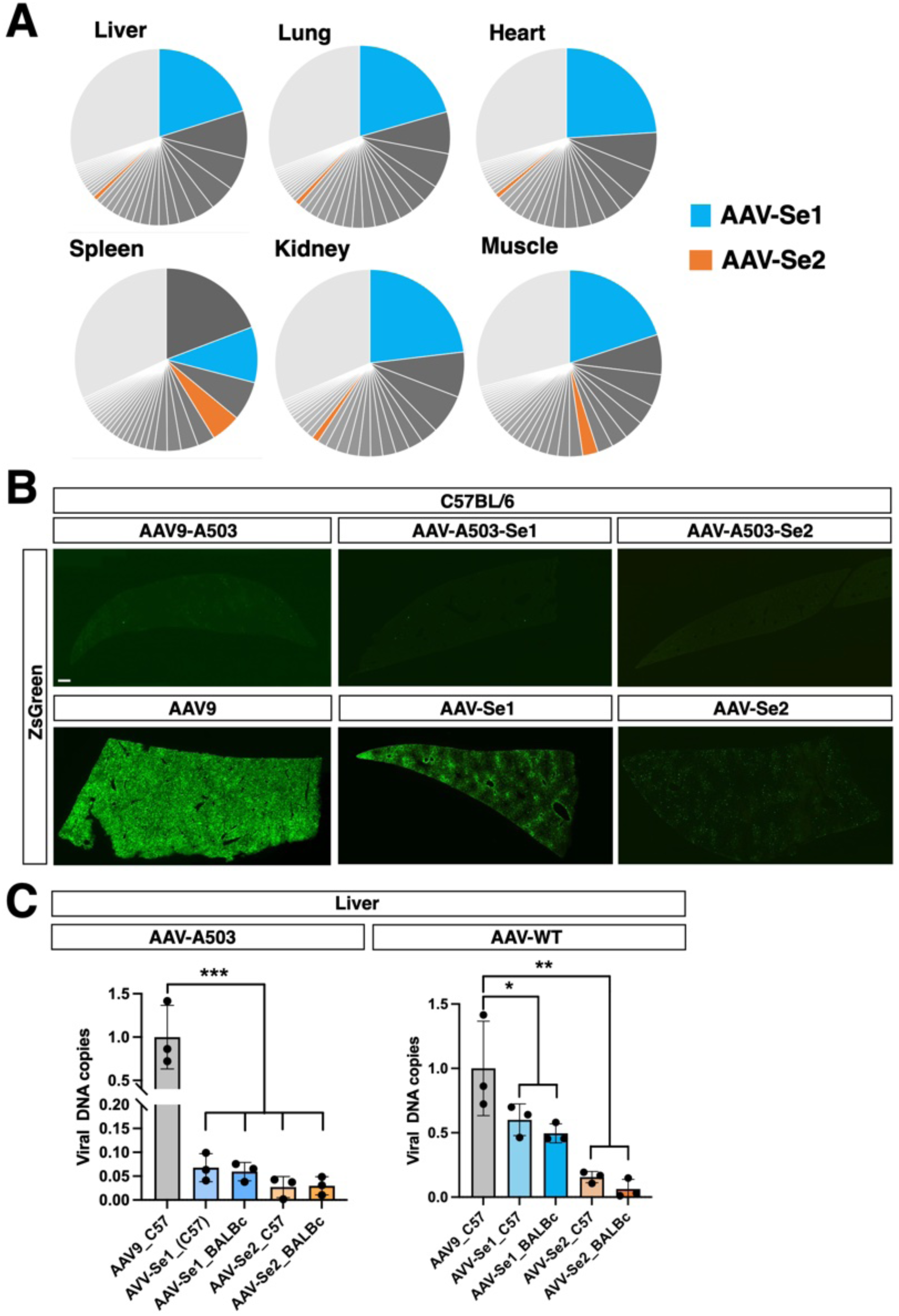
AAV-Se1/2 exhibit a lower tropism for liver with respect to the parental AAV9. **A)** Pie charts indicate the frequency of 7mer sequences determined by NGS in off-target organs, as in particular liver, lung, heart, spleen, kidney and muscle, at the fourth and last round of selection. In all off-target organs AAV-Se2 (orange) sequence was less enriched then AAV-Se1 (blue). **B**) Low magnification immunostaining for ZsGreen in livers derived from C57BL/6 mice treated with AAV9, AAV-Se1 and AAV-Se2 with (bottom panels) or without (A503, upper panels) competence for galactose binding. Scale bars, 200 µm. **C)** Viral DNA vector quantification by qRT-PCR in livers from C57BL/6 and BALBc mice treated with AAV9-A503, AAV-A503-Se1 and AAV-A503-Se2 (on the left) and AAV9, AAV-Se1 and AAV-Se2 (on the right). The results are reported as the fold change of viral ZsGreen DNA in mice treated with AAV-A503-Se1 and AAV-A503-Se2 (left) and AAV-Se1 and AAV-Se2 (right) relative to mice treated with AAV9 (n = 3 animals per group). Values are mean ± SD . *p < 0.05; **p < 0.01, ***p < 0.001. Statistical analysis is performed using one-way ANOVA followed by Tukey post-test.

### Analysis of AAV-Se1/2 brain tropisms in adult marmosets and human neurons

We next sought to determine whether AAV-Se1/2 conserved a higher brain transduction efficiency respect to the unmodified AAV9 in NHPs. To this end, a single dose (3 × 10^13^ vg) of AAV9, AAV-Se1 or -Se2, expressing the ZsGreen reporter, was injected through the femoral vein of adult marmosets previously selected for lacking any inherent immune response to AAV capsids. 3 weeks after viral administration the brains and peripheral organs were isolated and sections and lysates were prepared for immunohistochemistry and viral copy quantification, respectively (Figure 6A). During this time no adverse behavioral effects were observed in the treated animals with no gross changes in weight (Table S1). Additionally, liver enzyme levels and blood cell counts were normal at the time of sacrifice indicating the complete lack of enduring liver or systemic organic sufferance caused by either the surgery or the viral-based transgene expression (Table S2). ZsGreen staining on immunodecorated forebrain and cerebellar sections was homogeneously more intense after delivery with AAV-Se1 or -Se2 compared to AAV9 (Figure 6B). Likewise, the mRNA levels of the reporter were significantly higher respect to the AAV9 both in the cortex and cerebellum (2.2 ± 0.4 and 4.2 ± 0.9 for the cortex; 3.6 ± 0.8 and 4.7 ± 1.5 for the cerebellum for AAV-se1 and AAV-Se2, respectively) (Figure 6C). Of note, the numbers of cerebellar Purkinje cells infected with AAV-Se1/2 were significantly increased compared to that with AAV9 as revealed by Calbindin and ZsGreen co-staining (9% ± 3%, 14% ± 2%; 26% ± 5% of Calbindin^+^ Purkinje cells for AAV9, AAV-Se2 and AAV-Se1, respectively) (Figure 6D). Wide brain distribution of ZsGreen was visualized by DAB-mediated immunostaining in case of the AAV-Se2 showing overall homogeneous staining throughout the brain which was localized in cells with both neuronal, glial and endothelial morphology and with a relative enhanced transduction in the caudoputamen and thalamus (Figures 6E; S3A). Interestingly, quantification of DNA vector copy number indicated that AAV-Se1/2 compared to the parental AAV9 were endowed with higher transduction in the cerebral cortex and cerebellum, but had a significantly reduced tropism for the peripheral organs such as liver, muscle and heart (Figure S3B). To assess the cell-type specific transduction mediated by the AAV-Se1/2, we performed co-staining between the NeuN neuronal or Sox9 glial specific markers and the viral reporter ZsGreen. Interestingly, both AAV-Se1/2 sustained significantly higher transductions in neurons and astrocytes compared to AAV9 on cortical sections of the infected brains (AAV9: 1% ± 0.4% MAP2^+^ neurons, 3% ± 1% Sox9^+^ astrocytes; AAV-Se1: 2.5% ± 0.8% MAP2^+^ neurons, 7% ± 2% Sox9^+^ astrocytes; AAV-Se2: 6% ± 2% MAP2^+^ neurons, 9% ± 3% Sox9^+^ astrocytes). (Figures S2B,C).

**Figure 6:**
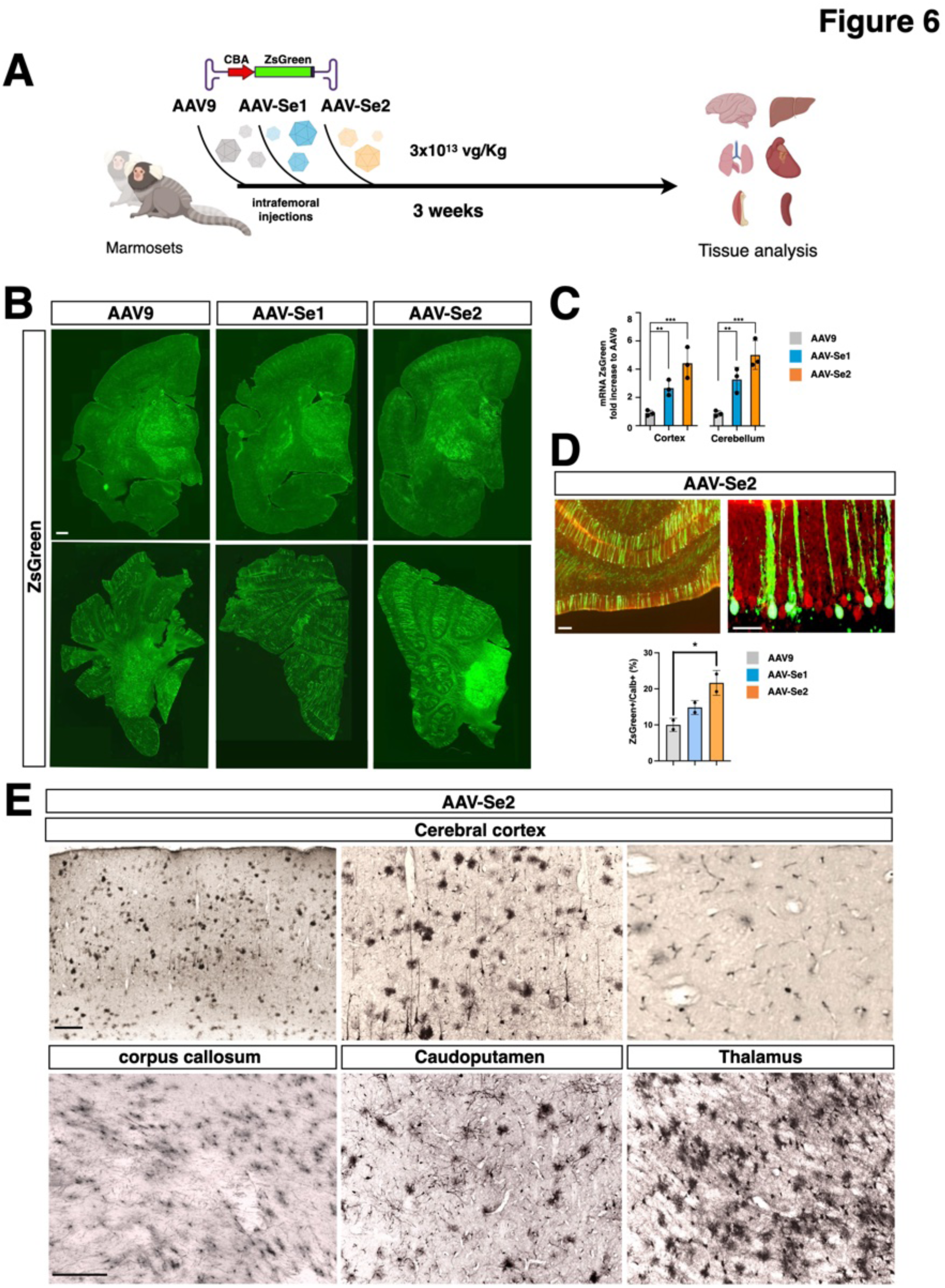
Brain transduction efficiency of AAV9 and AAV-Se1/2 in adult marmosets. **A)** Illustration of the experimental setting. Adult marmosets were systemically injected with AAV9 (n=1), AAV-Se1 (n=1) and AAV-Se2 (n=2) carrying the ZsGreen transgene under the control of the CBA strong constitutive promoter (experimental viral dose: 3 × 10^13^ vg/kg). After three weeks from the injections, marmosets were sacrificed for tissues analyses. **B)** Low magnification images of ZsGreen immunofluorescence in cerebrum (upper panel) and cerebellum (bottom panel) derived from marmoset treated with AAV9, AAV-Se1 and AAV-Se2. Scale bars, 200 µm. **C)** Bar graphs depicting fold change differences of ZsGreen mRNA levels in cortices and cerebella transduced with AAV-Se1/Se2 respect to AAV9 (n= 3 pieces for each tissue) . **D)** Low (upper) and high (bottom) magnification images of immunostainings for ZsGreen and Calbindin (Calb) in the cerebellum derived from marmoset treated with AAV-Se2. Scale bars, 100 µm. Below, quantitative assessment of transduced (ZsGreen^+^) Purkinje cells (Calb^+^) in brain derived from AAV9, AAV-Se1 and AAV-Se2 treated marmosets (n= 3 different fields of each cerebellum). **E)** Representative images of immunohistochemistry for the reporter ZsGreen expressed by the AAV-Se2 in in selected brain regions such as cerebral cortex, corpus callosum, caudoputamen and thalamus. Values are mean ± SD. *p < 0.05; **p < 0.01, ***p < 0.001. Statistical analysis is performed using one-way ANOVA followed by Tukey post-test.

To determine whether AAV-Se1/2 can also transduce human neurons, we tested them on human neuronal cultures differentiated from induced pluripotent stem cells (iPSCs). 6-weeks old iPSC-derived cortical neuronal cultures were infected with ZsGreen expressing AAV-Se1/2 or relative control viruses including PHP.eB, wild-type and A503 mutant AAV9 (Figure 7A). Interestingly, AAV-Se1/2 and PHP.eB exhibited comparable levels of infectivity of MAP2^+^ neurons that were significantly higher to that of wild-type AAV9 (AAV-Se2 41% ± 4%, AAV-Se1 36% ± 3%, PHP.eB 29% ± 4%, AAV9 21% ± 3%) (Figure 7A). In contrast the mutant AAV9 failed to support gene transfer in human neurons indicating that galactose binding is an absolute requirement for the AAV9 capsid to transduce neurons *in vitro*.

**Figure 7:**
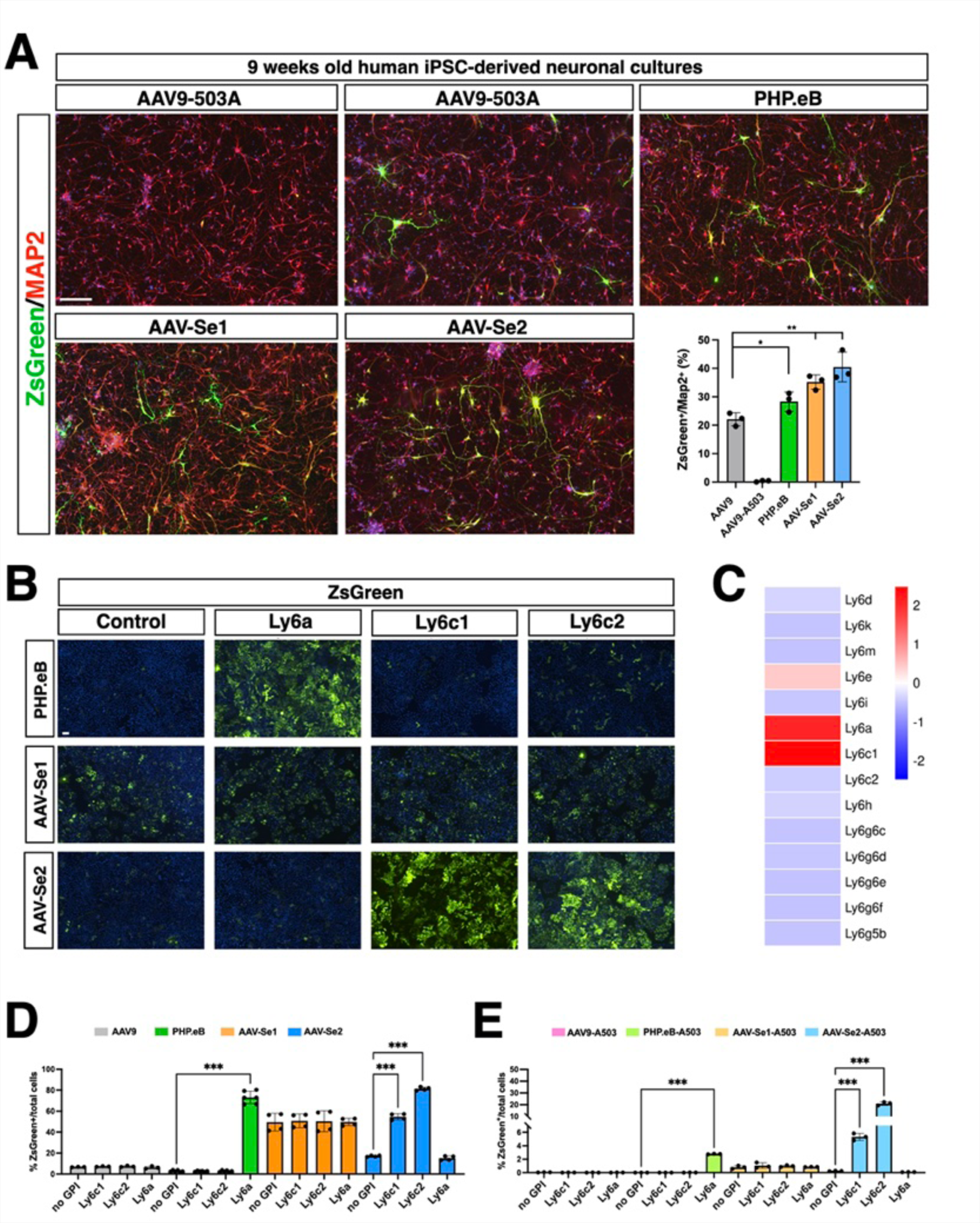
Viral transduction efficiency in human neurons and testing of Ly6 proteins as candidate receptors for AAV-Se1/2 capsids. **A)** Immunostaining for ZsGreen and MAP2 in hiPSCs-derived neurons transduced with AAV9-A503, AAV9, PHP.eB, AAV-Se1 or AAV-Se2 viral vectors. Bottom right, quantification of ZsGreen^+^ on total number of MAP2^+^ (n= 3 independent experiments). Scale bar, 100 µm. **B**) ZsGreen immunofluorescence in HeLa cells transfected with Ly6a, Ly6c1, Ly6c2 or untrasfected (control) and subsequently transduced with ZsGreen expressing PHP.eB, AAV-Se1 or AAV-Se2 viral vectors. Scale bars, 100 µm. **C**) Relative expression levels of Ly6 genes in brain endothelial cells as elaborated from a publicly available scRNA-seq dataset (https://doi.org/10.22002/D1.2090). **D,E**) Quantification by flow cytometry of ZsGreen positive HeLa cells transfected with Ly6a, Ly6c1, Ly6c2 or untrasfected (control) and subsequently transduced with ZsGreen expressing AAV9, PHP.eB, AAV-Se1 or AAV-Se2 viral vectors (**D**) or AAV9-A503, PHP.eB-503, AAV-Se1-A503 and AAV-Se2-A503 viral vectors (**E**). Values are mean ± SD of n = 3 independent experiments. *p < 0.05; **p < 0.01, ***p < 0.001. Statistical analysis is performed using one-way ANOVA followed by Tukey post-test.

### AAV-Se1/2 viruses showed different dependance by the LY6c1/2 receptors

Next, we sought to confirm that the AAV-Se1/2-mediated transduction was independent by the PHP.B receptor Ly6A. For this goal, we expressed Ly6a in HeLa cells, since giving their human origin they do not express it endogenously and assessed whether this *in vitro* setting was sufficient to discriminate the PHP.eB dependence by Ly6A. In fact, PHP.eB showed a marked increase in infectivity only when HeLa cells expressed its receptor Ly6A (Figure 7B). Encouraged by this result, AAV-Se1 and -Se2 were used to infect HeLa cells expressing or not Ly6a, but both of them resulted poorly susceptible to AAV-Se1/Se2 suggesting that Ly6A did not confer any advantage for AAV-Se1/Se2 transduction (Figure 7B). Ly6A is one member of the Ly6 family of GPI-anchored proteins highly expressed in the endothelium^28–31^. However, among the different Ly6 homologues, the Ly6C1 protein, share a similar enrichment in brain endothelial cells (Figure 7C). Thus, we sought to test whether AAV-Se1 and -Se2 showed any dependance on Ly6C1 or its highly homologue protein Ly6C2 for cell transduction. Remarkably, while gene transfer efficiency was not changed with AAV-Se1, we scored a significant increase in AAV-Se2-mediated expression in both Ly6C1- and Ly6C2-expressing cells. These results clearly suggested that AAV-Se2, but not AAV-Se1, infectivity is at least partly mediated by both LY6C1 and LYC2 receptors. Moreover, we quantified transduction efficiency by ZsGreen fluorescence flow-cytometry (Figures 7D, S4). From this analysis, it is noticeable that basal PHP.eB infectivity was minimal in HeLa cells not expressing any GPI protein (2.8% of total cells), as well as AAV9 (6.7% of total cells) whereas it was much higher for AAV-Se2 (17.2% of total cells) and even more for AAV-Se1 (49.5% of total cells). Thus, some levels of infectivity of the AAV-Se capsids can occur in HeLa cells in basal conditions, suggesting additional viral/cell interactions which sustain successful gene transfer. Ly6E is another homolog receptor of the family with high expression in brain endothelial cells and conserved in all mammalian species^32, 33^. Thus, we generated LY6E deficient HeLa cells by introducing a homozygous indel mutation by CRISPR/Cas9 technology (Figure S5). Interestingly, AAV-Se2, but not AAV-Se1, showed a reduced infectivity in Ly6E mutant HeLa cells (Figure S5). Thus, these results suggest that successful AAV-Se2 transduction is mainly dependent by the LYC1/2 receptors, but LY6E can yet sustain a residual infectivity of this AAV variant possibly acting as an alternative low-affinity receptor. Moreover, the presence of the W503A mutation was detrimental for cell transduction that dropped of 130, 62 and 70 times for AAV9, AAV-Se1 and AAV-Se2, respectively. No ZsGreen^+^ cells were detectable in HeLa cells infected with PHP.eB-A503 (Figure S4). Interestingly the mutant counterpart maintained, although reduced, their dependency on specific GPI protein (Figure 7E). Therefore, HeLa cells expressing Ly6a were infected significantly more by PHP.eB-A503 compared to control cells. Similarly, AAV-A503-Se2 transduced more HeLa cells that expressed LY6C1 and to greater extent LY6C2. AAV9-503 and AAV-A503-Se1 infection was not perturbed by the Ly6 GPI proteins exploited in this analysis.

### AAV-BI30 with mutated galactose binding domain significantly reduces liver targeting without affecting its brain transduction efficiency

Given that the loss of galactose binding on the AAV-Se1/2 dramatically reduced their off-targeting to the liver while mostly preserving their transduction efficiency for brain endothelial cells, we thought that the same approach could be extended to other endothelial-directed capsids. In particular, the AAV9 engineered capsid variant AAV-BI30 was recently showed to efficiently transduce the endothelial cells of the brain, retina and spinal cord vasculature after intrasystemic delivery^34^. However, AAV-BI30 also exhibited an extensive tropism for peripheral organs, and in particular for the liver, which could lead to hepatotoxicity in the treated mice^34^. Thus, in analogy to our previous results, we introduced the W503A mutation within the AAV-BI30 capsid to assess its impact of the infectivity profile of this capsid. As expected, the ZsGreen expressing AAV-A503-BI30 intravenously administrated the at 1 × 10^11^ vg per mouse, sustained an efficient gene transfer in the brain vasculature, but associated with a strong reduced tropism for liver respect to its unmodified parental capsid (Figure 8). Thus, modulating the galactose binding ability of the capsid variants can be an additional change that has great impact on restricting the off-target viral infectivity in peripheral organs. In contrast, this variation will not markedly affect the tropism-of-interest if this is mainly mediated by the engineered modification inserted in the capsid variant.

**Figure 8:**
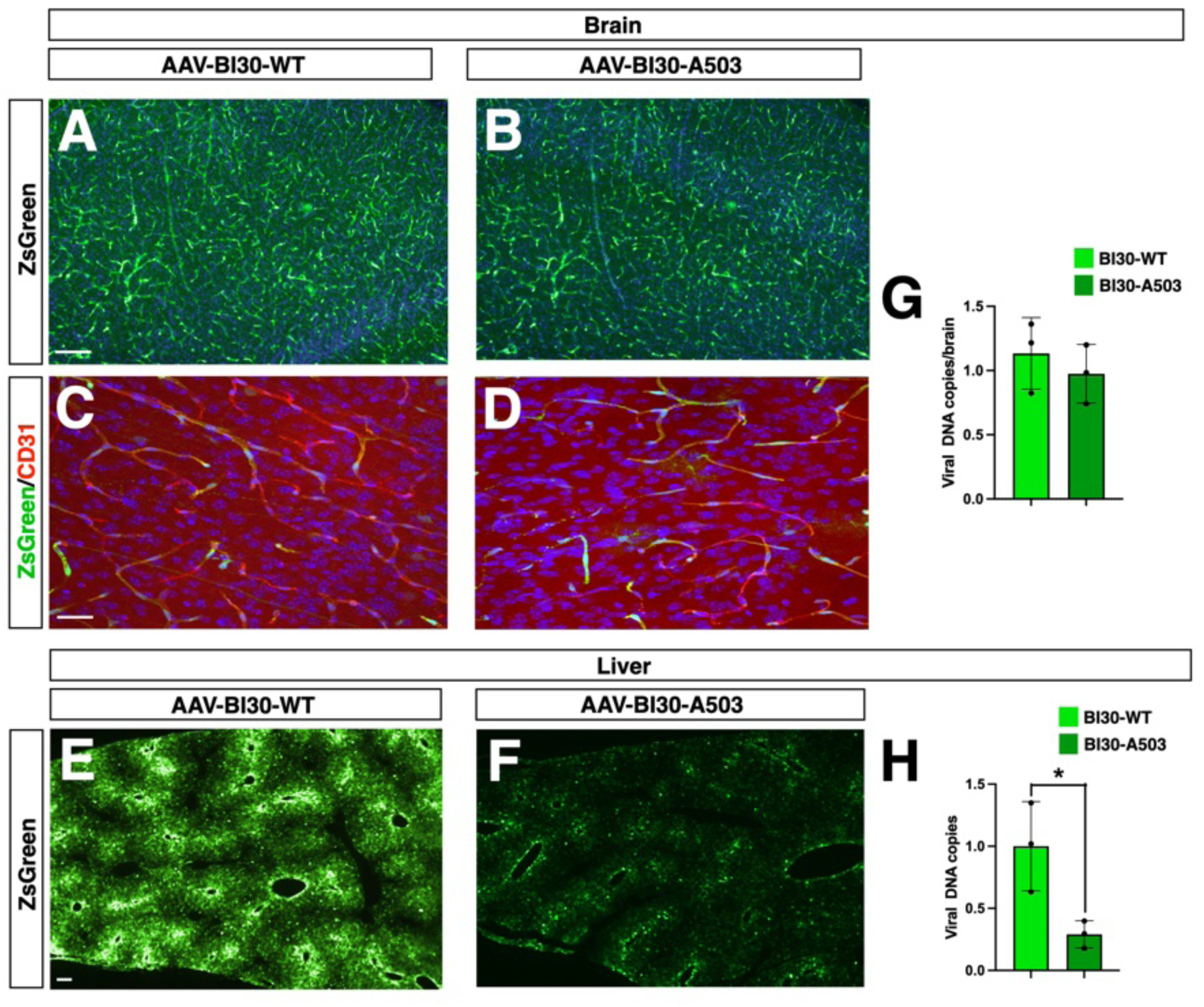
Effective liver detargeting by inserting the A503 mutation in the AAV-BI30 capsid. **A-D)** Single ZsGreen (**A**,**B**) or double ZsGreen/CD31 (**C**,**D**) immunofluorescence staining highlighting the enriched endothelial the transduction of the unmodified AAV-BI30 (**A**,**C**) and its mutant form AAV-BI30-A503 (**B**,**D**) capsid. Scale bars, 100 µm. **E,F)** Representative images of ZsGreen immunefluorescence levels in liver of mice intravenously injected with the unmodified AAV-BI30 (**E**) and its mutant form AAV-BI30-A503 (**F**) capsid. Scale bar, 200 µm. **G,H)** Viral DNA copies quantification by qPCR in brain (**G**) and liver (**H**) tissues transduced with either unmodified AAV-BI30 or its mutant form AAV-BI30-A503 capsid. Values are mean ± SEM of n = 3 independent experiments. ***p < 0.001. Statistical analysis is performed using unpaired *t*-test.

## Discussion

Herein, we described the isolation of two novel AAV9 engineered variants with a superior ability to target the brain after intravenous delivery by screening a new display peptide library developed from a galactose binding-deficient AAV9 capsid. We speculated that this combination would erase the natural binding pattern of the mutant capsid maximizing the selection of new capsid-cellular receptor interactions exquisitely depending by the exogenous peptide. However, although the AAV-A503-Se1/2 viruses showed an efficient transduction of the brain vasculature, these capsids exhibit a very poor capacity to infect brain cells. Only after restoring the functional galactose binding site, both viruses acquired the ability to infect both neuronal and glial cells in the brain parenchyma. These results indicate that AAV9-like capsids might require two distinct domains to efficiently transduce the brain: the first required to bind and cross the endothelium which is provided by the exogenous peptide and the galactose binding to subsequently transduce the other brain cells. No case has yet been described in which a single domain in the capsid can sustain both functions of the same virus. On this light, our approach was well suited for isolating peptides that could confer to the capsids an enhanced transduction of the endothelium, but without directly influencing the subsequent tropism toward either neuronal or glial cells which is mainly controlled through the virus-galactose interaction.

While AAV-Se1 emerged already at the second cycle, the AAV-Se2, was found enriched only at the fourth and last selection round suggesting that the dynamics of enrichment could be extremely different between the different capsid variants. This is reflected by the rather different peptide sequence selected in the two capsids and their likely distinct cell receptors. Both sequences are distinct from those of other previously selected variants, suggesting that the diversity offered by the randomized heptapeptide is extremely high making unlikely the selection of the same exact sequences in independent experiments. The AAV-A503-Se1/2 viruses showed a very selective transduction of brain endothelial cells both in C57BL/6 and BALBc mice strains with a very minority of non-endothelial cells to be infected. Furthermore, AAV-A503-Se1/2 exhibited an extremely low tropism compared to the parental AAV9 for peripheral organs, including the liver, thereby, avoiding risks of systemic toxicity or off-targets effects even when relatively high viral doses are administrated intravenously. Thus, these viruses are precious tools for the selective and efficient gene transfer within the brain vasculature with superior advantages respect to previous similar capsids. Conversely, after galactose-binding reconstitution the wild-type AAV-Se2, more than the AAV-Se1, acquired a strong efficiency to transduce both neuronal and glial cells across the entire brain. In fact, AAV-Se2 transduction levels in the brain tissue were not significantly different from those sustained by the PHP.eB, one of the capsids with the highest brain targeting after intravenous delivery^27^. Even in this case, anyhow, wild-type AAV-Se1/2 displayed a significantly lower tropism for the liver compared to the unmodified AAV9 suggesting that the integrated peptides have a direct role in de-targeting AAV-Se1/2 capsids from this peripheral organ. We, next, evaluated the transduced rate of AAV-Se1/2 in the brains of adult marmosets. In a side-by-side comparison, we confirmed that these variants have a higher brain tropism respect to the unmodified AAV9. In particular, cerebellar Purkinje cells were efficiently transduced by the AAV-Se2 with an overall rate superior to 25%. Thus, this capsid represents an interesting tool for gene delivery to treat genetic forms of ataxia caused by selective degeneration of this class of neurons. Although we could work with only few animals given the difficulty to access them during the COVID-19 pandemic period, we treated two animals at least with the AAV-Se2 capsid obtaining very comparable data and, thus, indicating a general reliability of the technical procedures uniformly applied to all the animals. In overall, the infectivity of both AAV-Se1/2 in marmosets was generally lower compared to that detected in mice indicating a lack of conserved gene transfer efficiency in NHPs. For this reason, we considered to evaluate the dependance of these viruses on receptors already implicated in the virus-cellular interactions. At first, we set up an in vitro assay to determine the efficiency of viral transduction on HeLa cells expressing a candidate viral receptor. To validate the system, we showed that only cells expressing the receptor Ly6a are robustly transduced by the PHP.eB virus. Ly6A belongs to a large super-family of small proteins with a LU domain composed of 80 amino acids containing ten cysteines which creates a three-fingers spatial configuration^35, 36^. Despite the mouse genome contains 61 Ly6 members, Ly6A shares the highest homology with proteins encoded by adjacent genes forming the so-called Ly6A sub-family within the same genomic locus, which has been largely deleted during mammalian evolution^37^. Surveying the expression profile from publicly available databases, we found that among these homologues Ly6C1 presents the highest and most selective expression in brain endothelial cells similarly to Ly6A. Remarkably, we found that transduction of AAV-Se2 was strongly enhanced in cells expressing either Ly6c1 or its close homolog Ly6c2, suggesting that they are plausible receptors for this capsid. Moreover, we also found that AAV-Se2 infectivity can be augmented in cells expressing Ly6e. Thus, the enhanced brain transduction of AAV-Se2 respect to AAV9 in NHPs is likely mediated by Ly6E which, contrary to Ly6c1/2, is conserved throughout mammalian evolution. Moreover, given that endothelial expression of Ly6c1 and Ly6e is mostly confined to the brain, this would contextualize the very poor tropism of AAV-Se2 that we observed in the peripheral organs both in mice and NHPs. In contrast, we noted that the basal levels of transduction in cells not expressing any murine Ly6 GPI proteins were significantly higher for AAV-Se1. Thus, this capsid has acquired other properties that have positively influenced its infectivity that might explain its high and selective transduction of the brain endothelium.

Our study clearly showed that viral tropism can be modulated in a combinatorial fashion by altering the galactose-binding domain in AAV9 engineered capsids. This option has remained poorly exploited in the field while we have shown that it enables to minimize or totally prevent the tropism for liver and other peripheral organs, while preserving the infectivity toward the tissue targeted through the displayed peptide. We provide evidence that this approach can be generalized to other viral vectors by showing that the strong liver tropism of the AAV-BI30 capsid can be significantly dampened, although not reaching the minimal levels featured by the AAV-A503-Se1/2 variants, while keeping unaltered its infectivity of the brain endothelium. We postulate that the same modification might work with several other engineered capsids to enhance their safety profile. Thus, this option further expands the possible targeted manipulations on the capsid variants to control their viral transduction pattern and significantly minimize off-targeting tropism, that are indispensable features to nominate them for clinical exploitation.

## Acknowledgements

We thank D. Bonanomi and L. Naldini for providing valuable reagents as well as T. Sieber and M. Trepel for generating and providing the p9K7 plasmid library. We are grateful to all members of the Broccoli’s lab for helpful discussion. We acknowledge the FRACTAL and ALEMBIC core facilities for expert supervision in flow-cytometry and confocal imaging, respectively. This work was supported by Telethon Foundation (#1350) and PNRR-National Center for Gene therapy and drugs based on RNA technology (CN00000041).

## Author contributions

S.S.G. and M.L. performed all the experiments and analyzed data; B.B. contributed to gene cloning and qPCR quantifications; A.I. analyzed viral infections in iPSC-derived neuronal cultures; I.P. supervised the work with marmosets; J.K. supervised the plasmid and viral library generation; V.B. supervised, coordinated and supported the project and wrote the manuscript.

## Competing interests

The authors have filed a patent application for the work described in this manuscript.

## Methods

### Mice

Mice were maintained at San Raffaele Scientific Institute Institutional mouse facility (Milan, Italy) in micro-isolators under sterile conditions and supplied with autoclaved food and water. C57BL/6 and BALBc mice were purchased from Charles River Laboratories Italia (Calco, Italy) and were maintained in their background. All procedures were performed according to protocols approved by the internal IACUC and reported to the Italian Ministry of Health according to the European Communities Council Directive 2010/63/EU.

### Marmoset monkeys

Four adult (3-5y) male common marmosets (*Callithrix jacchus*), with a body weight between 280-330 g, were purchased from the purpose-bred colony of the Biomedical Primate Research Centre (BPRC) in the Netherlands. Prior to inclusion in the study, the monkeys underwent elaborate physical examination by the institute’s veterinarian, as only healthy monkeys were included. All animals were experimental naïve and in good health and free of pathogenic ecto- and endoparasites and common bacteriological infections: Yersinia pestis, Yersinia pseudotuberculosis, Yersinia enterocolitica, Salmonella, Shigella, Aeromonas hydrophilia. They also showed a negative tuberculin reaction. Before the selection of the monkeys a blood sample was taken and tested for lacking any inherent immune response to AAV capsids. The monkeys were pair-housed under conventional conditions in spacious cages with a varying cage environment and were under intensive veterinary care throughout the study. All monkeys were observed for clinical symptoms and the body weight was measured every week. The monkey facility was under controlled conditions of humidity (>60 %), temperature (22–6 °C) and lighting (12 h light/dark cycles from 7 am to 7 pm). The animals were daily fed with standard monkey-chow (Special Duit Services, Witham, Essex, UK) in combination with fruits and vegetables and ad libitum water supply. This study was performed under project license AVD5020020172687, which was issued by the competent national authorities (Central Committee for Animal Experiments) according to Dutch law, article 10a of the *“Wet op de Dierproeven”*. Approval was obtained by the institutional animal welfare body (CCD 008B). All procedures, husbandry, and housing were performed in accordance with the Dutch laws on animal experimentation and the EU Directive 63/2010. The BPRC is accredited by *AAALAC International*.

### Preparation of the random AAV9 display peptide library

A random AAV9 display library displaying heptapeptide insertions at amino acid position A589 (VP1 numbering) with a diversity of 1 × 10^8^ unique clones (calculated at plasmid level) was produced using a three-step protocol as described previously^13, 38, 39^. In short, a degenerate oligonucleotide encoding seven random amino acids (encoded by NNK to avoid two out of three stop codons and to limit the number of degenerate codons) was synthesized commercially (Metabion) as follows: 5’-AGAGTGGCCAAGCAGGC(NNK)7GCCCAGGCGGCCACCGGC-3’. Second-strand synthesis was performed using the Sequenase enzyme (Affymetrix) and the primer 5’-GCCGGTGGCCGCCT-3’. The double-stranded oligonucleotide insert was cleaved with BglI, purified with the QIAquick Nucleotide Removal Kit (Qiagen), and ligated at nucleotide position 3967 of the AAV genome into the SfiI-digested p9K7 library plasmid carrying a W to A mutation at position 503 (VP1 numbering). Electrocompetent DH5a bacteria were transformed with the library plasmids using the Gene Pulser Electroporation System (Bio-Rad). The diversity of the plasmid library was determined by the number of clones growing from a representative aliquot of the transformed bacteria on LB agar containing 150 mg/ml ampicillin. Library plasmids were harvested from transformed bacteria and purified using the NucleoBond Xtra MIDI plasmid preparation kit (Macherey-Nagel). The AAV library genomes were packaged into 293T cells by transfecting 1 × 10^7^ in ten 15-cm with the library plasmids (containing the cap gene with peptide insertion, with inverted terminal repeats as packaging signal)^13^, and the pXX6 adenoviral helper plasmid. Transfection was performed using Polyfect (Qiagen) according to manufacturer instructions. The final random peptide AAV display library was harvested approximately 72 h after transfection both by lysis of the harvested cells (hypertonic shock) and precipitation and concentration of particles present in the supernatant (using 8% polyethyleneglicol 8000). Thus, library particles were purified by iodixanol density gradient by ultracentrifugation^40^. A discontinuous iodixanol gradient was prepared by subsequently underlying the harvested library particles with 15, 25, 40, and 40% iodixanol solutions, followed by ultracentrifugation at 190,000 g for 180 min. The 40% iodixanol fraction, containing purified viral particles, was aspirated and concentrated using Amicon15 columns (MERK-Millipore). The virus titer was determined using AAVPro titration kit (Takara).

### In vivo screening of the random AAV9 display peptide library

For the in vivo selection, 1 × 10^11^ genomic library particles were injected into the tail vein of BALBc mice. A week later, mice were sacrificed and the complete brain as organ of interest was removed. Total tissue DNA was extracted using the DNeasy Tissue Kit (Qiagen). The random oligonucleotide insertions from the enriched AAV library particles were amplified by PCR using the primers 5’–GGAGCTTCTTCTTGGGCTCT–3’ and 5’– AGCGGAGAAGGGTGAAAGTT–3’. Approximately 20 PCRs with lg template DNA each were set up. The amplicon was purified Wizard SV Gel and PCR Clean-up System (Promega) and was used to produce preselected libraries, using the same procedure described above for the random peptide display library, beginning from SfiI digestion. This procedure was applied to subsequent rounds of selection. Four rounds of selection were performed in n = 1 animal each. After each round of selection, ten clones were sequenced. Selected library clones were produced as recombinant AAV vectors for further analysis. In the fourth round of selection not only the on-target tissue but also several off-target organs (liver, spleen, lung, kidney, heart, and muscle) were collected for DNA extraction and PCR extraction as described above.

### Next-Generation Sequencing

Amplicon libraries for Illumina MiSeq next-generation sequencing were amplified for each of four preselected libraries and for all off-target organ of the fourth selection. 1ng of the amplicon described above was used as template in a PCR reactions using the following pairs of linker primers fw: 5’-<u>ACACTCTTTCCCTACACGACGCTCTTCCGATCT</u>CAGCCACAAAGAAGGAGAGG-3’and rv: 5’-<u>GACTGGAGTTCAGACGTGTGCTCTTCCGATCT</u>GTCCTGCCAAACCATACCC -3’ Which include illumine adapters (underlined sequences). The amplicon was purified submitted at Azenta for Amplicon EZ sequencing.

An analytical gel electrophoresis was performed with parts of each probe to determine the concentration of the corresponding fragments, the probes were diluted and 1 ng (5 × 10^9^ copies) of each probe were used as templated for a second PCR with the NGS forward primer: NGS2_fwd: 5′-AATGATACGGCGACCACCGAGATCTACACTCTTTCCCTACACGAC-3′ in combination with one of the individual Illumina index adapters: 5′-CAAGCAGAAGACGGCATACGAGAT(7nt-index)GTGACTGGAGTTCAGACGT GTG-3′. Each sample was sequenced with a depth of at least 500 000–reads. Data analysis was performed by a custom script.

### Computational analysis

FASTQ reads were quality checked and trimmed. Random oligonucleotide sequences were identified by searching sequence reads for exact matches of the two sequences CCAAGCAGGC and GCCCAGGCGG, which flank the random insert within the AAV9 cap sequence. The obtained sequences were sorted into clusters of identical sequences and subsequently different filters were applied to remove possible artifacts. First, sequences whose length diverged from the expected length were not considered for further analysis. To remove sequences containing sequencing errors, it was then checked for every sequence if it was at least 100 times less abundant than any other sequence. If the Levenshtein distance between two such sequences was 1, we excluded the less abundant sequence from further analysis. The remaining sequences were translated into peptides. During this process, only sequences were counted that were composed of nucleotide trimers matching the coding scheme of the oligonucleotide synthesis (Figure 1). Downstream statistics and Plot drawing were performed with R script.

### Generation of packaging plasmids and gene transfer vectors

Plasmid encoding the modified AAV capsid of interest were subcloned into tTA-iCAP-PHP.eB (Addgene). The mutated version of Se1 and Se2 were generated using as donor sequence the clones derived from the third round of selection which contained the heptapeptide at position 589 and the W503A mutation (VP1 numbering). The restriction fragment SapI-AfeI purified from the Se1 and Se2 clones was cloned into the vector tTA-iCAP-PHP.B digested NheI-AfeI and ligated with a second restriction fragment (NheI-SapI) extracted from the same vector, thus originating the plasmids for the production of AAV-Se1-A503 and AAV-Se2-A503 vectors. Conversely wild-type plasmids coding for W in position 503 of VP1 (AAV-Se1, AAV-Se2 and AAV9) were produced using PCR amplification of the CAP cistron of tTA-iCAP-PHP.eB using a common fw primer 5’-ACC**GCTAGC**TTCGATCAACTACGCG-3’ and reverse primers containing the desired heptapeptide ant position 589 (underlined) or the native sequence of AAV9 flancked by an AgeI site (bold):

Se1-Rv:

5’CCCA**ACCGGT**GGCCGCCTGGGC<u>ACCCACAGAACGAACACCATT</u>GCCTGCTTGGCCACTCTGGTGGT TTGTGGCCACTTGTC -3’;

Se2-Rv:

5’CCCA**ACCGGT**GGCCGCCTGGGC<u>AAACCGCCCCGGAACACCAGG</u>GCCTGCTTGGCCACTCTGGTGGT TTGTGGCCACTTGTC-3’;

AAV9-Rv:

5’-CCAA**CCGGTC**TGCGCCTGTGCTTGGGCACTCTGGTGGTTTGTGGCCACTTGTC-3’.

For AAV9-A503 the same oligos for AAV9 were used but as template AAV-Se2-A503 was used. Amplicons were restriction digested with NheI and AgeI as well as tTA-iCAP-PHP.eB vector and cloned. The coding sequences for ZsGreen (pIRES2-ZsGreen1, Takara) and Luciferase (*luc2*, pGL4.14-Glo, Promega) were cloned into a ssAAV-CBA vector (Luoni et al., 2020).

### AAV Virus production and purification

AAV replication-incompetent, recombinant viral particles were produced 293T cells, by polyethylenimine (PEI) co-transfection of three different plasmids: the gene transfer vector, the packaging plasmid (see above) and pHelper (Agilent) for the three adenoviral helper genes. The cells and supernatant were harvested at 120 hrs. Cells were lysed in hypertonic buffer (40 mM Tris, 500 mM NaCl, 2 mM MgCl_2_, pH=8) containing 100U/ml Salt Active Nuclease (SAN, Arcticzymes) for 1h at 37°C, whereas the viral particles present in the supernatant were concentrated with 8% PEG8000 (Polyethylene glycol 8000, Sigma-Aldrich) by precipitation (4000g, 20min). In order to clarify the lysate cellular debris where separated by centrifugation (4000g, 30min). The viral phase was isolated by iodixanol step gradient (15%, 25%, 40%, 60% Optiprep, Sigma-Aldrich) in the 40% fraction and concentrated in PBS (Phosphate Buffer Saline) with 100K cut-off concentrator (Amicon Ultra15, MERCK-Millipore). Virus titers were determined using AAVpro Titration Kit Ver2 (TaKaRa).

### Lentivirus Virus production and purification

GPI coding sequences (Ly6a: NM_001271416; Ly6c1: NM_001252055; Ly6c2: NM_001099217) were gene synthetized (Azenta) with flanking restriction sites apt for sub-cloning into the LV-Ef1a backbone^41^. Lentiviral replication-incompetent, VSVg-coated lentiviral particles were packaged in 293T cells^42^. Cells were transfected with 30μg of vector and packaging constructs, according to a conventional CaCl_2_ transfection protocol. After 30 h, medium was collected, filtered through 0.44μm cellulose acetate and centrifuged at 20000 rpm for 2 h at 20°C in order to concentrate the virus.

### In vivo administration of AAV vectors in mice and tissue collection

Vascular injection was performed in a restrainer that positioned the tail in a heated groove. The tail was swabbed with alcohol and then injected intravenously with a viral concentration (1 × 10^12^ to vg/mL) in a total volume of 100 µl of recombinant AAV particles in PBS. C57BL/6 and BALBc mice, independently of sex, were randomized in groups and injected in the tail vein between 30 and 40 days of age. Following injection, all mice were monitored for adverse effect and two weeks after infection were sacrificed and tissues harvested. Briefly, mice were anesthetized with ketamine/xylazine and transcardially perfused with 0.1 M phosphate buffer (PB) at room temperature (RT) at pH 7.4. Upon this treatment brain and liver were collected. Brain hemispheres were separated: one half was post-fixed in 4% paraformaldehyde (PFA) for two days and then soaked in cryoprotective solution (30% sucrose in PBS) for immunofluorescence analysis, the other half was further sectioned in different areas (cortex and cerebellum) quick frozen on dry-ice for Western blot, RNA and DNA extraction. Liver specimens were collected similarly.

### Testing basal immunity to AAV capsids in NHPs

Sera of six untreated animals were collected for neutralizing antibodies detection: Hela cells were plated in 96-well plate (6 × 10^4^ cells/cm^2^). The day after Monkey sera and Pooled Human Sera (PHS, Dunn Labortechnik GmbH) were heat inactivated (56°C, 30’), filtered (0,22 µm) and serially diluted in semi-log fashion. Each dilution was supplemented with AAV9-Luciferase (1 × 10^8^ vg/well) and negative (no virus) and positive controls (no serum) were sat. After incubation (1h, 37°C) these solutions were added to Hela Cells and incubated overnight (37°C, 5% CO_2_). The day after the plates were pre-cooled (RT, 15’) supplemented with luciferase substrate (Bright Glo Luciferase Assay System, Promega) incubated (RT, 30’) and luminescence was red with Victor^3^ microplate reader (Perkin-Elmer).

### In vivo administration of AAV vectors in marmosets and tissue collection

Clinical chemistry and hematological parameters were checked in blood samples for abnormalities before the start of the study, and for possible aversive effects of the treatment at the end of the study. AAVG, AAV-Se1 or –Se2 was administered intravenous (femoral vein) in a single dose (3 × 10^13^ vg) of in a volume of 1 ml under sedation with 12 mg/kg alfaxan IM (Vetoquinol, ‘s-Hertogenbosch, The Netherlands. After injection, the animals were closely monitored during a 3-hour recover period. Thereafter, the monkeys were daily observed for possible clinical signs like immobility, apathy, loss of appetite, and changes in movement. The body weight of the monkeys was measured once a week. During the study, the monkeys stayed in pairs together in their own home cage. Three weeks after the injection the monkeys were euthanized for collection of blood and tissues. Just before euthanasia, the monkeys were weighted and 2 ml serum for clinical chemistry, 2 ml plasma for hematology, and 2 ml EDTA blood for further analysis, was withdrawn from the femoral vein. Euthanasia was induced with Alfaxan 10 mg/kg + Ketamine 10 mg/kg IM. Immediately thereafter, the brains and other organs were removed in the section room. The whole brain was taken and weighted. The brain was cut into two hemispheres (including the brain stem). One hemisphere was stored in 4 % formalin for 48 hours. The other hemisphere was cut into different brain areas (cerebellum, pons, medulla, hippocampus, striatum, thalamus, temporal cortex, frontal cortex, visual cortex, putamen, and caudate nucleus) and snap frozen in liquid nitrogen and stored at -80° C. All the remaining parts of the brain were also stored at -80° C for qPCR analysis. From the other organs (kidney, liver, heart, spleen, lung, muscle hamstring, terminal ileum and colon) two parts were collected and stored in 4 % formalin (48 hours) or frozen in liquid nitrogen and stored at -80° C.

### Immunofluorescence

Tissues were sectioned using cryostat after optimal cutting temperature compound (OCT) embedding in dry ice. Free-floating 50μm-thick coronal sections were rinsed in PBS and were incubated with 10% donkey serum and 3% Triton X-100 for 1 hr at RT to saturate the unspecific binding site before the overnight incubation at 4°C with the primary antibody (diluted in the blocking solution). Fixed cells and tissues were washed with PBS (3×) and incubated with 10% donkey serum and Triton X-100 (3% for tissues and 1% for cells) for 1 hr at RT to saturate the unspecific binding site before the overnight incubation at 4°C with the primary antibody. Upon wash with PBS (3×), cells and tissues were incubated for 1 h at RT in blocking solution with DAPI and with Alexa Fluor-488 and Alexa Fluor-594 anti-rabbit, anti-mouse or anti-rat secondary antibodies (1:1000, ThermoFisher Scientific). After PBS washes (3×), cells and tissues were mounted with fluorescent mounting medium (Dako). Images were captured with a Nikon Eclipse 600 fluorescent microscope or with Leica TCS SP5 Laser Scanning Confocal microscope. Cell and tissue were stained with the following primary antibody: rabbit anti-ZsGreen (1:500; Takara), rabbit anti-NeuN (1:500; Merck), rabbit-Sox9 (1:500; Merck), rabbit-Calbindin (1:500; Swant), rat-CD-31 (1:500; BD biosciences), chicken-MAP2 (1:500; Abcam).

### Determination of vector copy numbers by qPCR

Total DNA was isolated from animal tissues (cortex, cerebellum, liver, heart, muscle) using the Qiagen DNeasy Blood & Tissue Kits (QIAGEN). The quantification of vector transgene expression was calculated by qRT-PCR relative to the ZsGreen transgene using the following primers: 5’-GCTCCTTCCTGTTCGAGGAC-3’and 5’-TCGGTCATCTTCTTCATCACG-3’. The DNA levels were normalized against an amplicon from a single-copy mouse gene: in murine samples: murine *Lmnb2*, amplified from genomic DNA using 5’-CTGAGGGTTGCAGGCAGTAG-3’and 5’-TGTGGACAGACCTGGGTAGG-3’; in simian samples: Callithrix jacchus LMNB2 amplified from genomic DNA using 5’-TTCTGGTCTCCATGCCACTC-3’and 5’-TGGTTCCTGTTGCGTCCTAC-3’. Depending on the control samples Normalization was applied.

### CRISPR/Cas9 gene editing in HeLa cells

We used the program CRISPOR (http://crispor.tefor.net) to design sgRNAs on exon 3 of the human LY6E gene (Gene ID: 4061). Efficiency of three different sgRNAs was compared in HEK293 cells and we selected sgRNA#1 (5’-agaaggcgtcaatgttggtg-3’) for its high activity of INDEL rate using the T7 Endonuclease I assay. Then, HeLa cells were co-transfected with the vectors U6-sgRNA#1-EF1α-Blast and the pCAG-Cas9-Puro using the Lipofectamine Transfection Reagent (Invitrogen)^43^. Co-transfected colonies were then selected by the combination of puromycin (1 μg/ml) and blastidicin (10 μg/ml) and then isolated through single colony picking. Finally, resistant cell clones were assessed by TIDE analysis followed by Sanger sequencing for isolating either clones with the targeted genomic modification in the LY6E gene or clones with the unmodified LY6E gene (control cells).

### Transduction assay in neuronal cultures and Hela cells

Primary neuronal cultures were prepared at embryonic day 17.5 (E17.5) from mouse embryos. Briefly, cortices were individually dissected, sequentially incubated in trypsin (0,005%, 20 min at 37°C) and DNAse (0,1mg/mL, 5min at room temperature) in HBSS (Hank’s buffered salt solution without Ca^2+^ and Mg^2+^). Cells wand plated on poly-L-lysine coated glasses (2 × 10^5^ cells/cm^2^) in Neurobasal medium enriched with 0,6% glucose, 0,2%penicillin/streptomycin, 0,25% L-glutamine and 1% B27. Viral particles were directly added to cultured neurons 3 days after seeding, with a final concentration 1 × 10^10^ vg/ml. iPSCs were initially differentiated in Neural Progenitors Cells (NPCs) as described in Iannielli et al.^44^. NPCs were, then, dissociated with Accutase and plated on matrigel-coated 6-well plates (3 × 10^5^ cells per well) in NPC medium. Two days after, the medium was changed with the differentiation medium containing Neurobasal, 1% Pen/Strep, 1% Glutamine, 1:50 B27, 10 μM SU5402, 8 μM PD0325901, and 10 μM DAPT was added and kept for 3 days. After 3 days, the cells were dissociated with Accutase and plated on poly-L-lysine/laminin/fibronectin (100 µg/ml, 2 µg/ml, 2 µg/ml)-coated 12-well plates (2 × 10^5^ cells per well) and 24-well plates (1 × 10^5^ cells per well) in neuronal maturation medium supplemented with ROCK inhibitor Y27632 (10 µM) for the first 24 h. Neuronal maturation medium was composed by Neurobasal, 1% Pen/Strep, 1% Glutamine, 1:50 B27, 20 ng/ml human BDNF, 200 μM Ascorbic Acid, 250 μM Dibutyryl cAMP, 10 μM DAPT, 1 μg/μl Laminin. The culture medium was replaced the next day to remove the ROCK inhibitor, and at this stage half of the medium was changed every 2–3 days. Viral particles were directly added to cultured neurons after six weeks of differentiation, with a final concentration 10^10^ vg/ml. All the analysis was conducted one week after the infection upon fixation of the specimen (4% PFA, 4°C, 30’).

HeLa cells were maintained in Dulbecco Modified Eagle Medium – high glucose containing 10% fetal bovine serum, 1% non-essential amino acids, 1% sodium pyruvate, 1% glutamine, and 1% penicillin/streptomycin. Cells were split every 3-4 days using Trypsin 0.25%. For transduction assay cells were plated (2 × 10^5^ cells/cm^2^) and infected at the first day *in vitro* (DIV1) with lentiviral vectors expressing GPI protein. At DIV 3 cells were replated (2 × 10^5^ cells/cm^2^) and infected with different AAV recombinant virus carrying ZsGreen transgene (4 × 10^9^ vg/ml) and finally at DIV6 cells were fixed for immunofluorescence staining (4% PFA, 4°C, 30’) or for FACS analysis (4% PFA, 4°C, 10’). In the former case cell were washed, resuspended (PBS, 2% FBS) and filtered (70 mm), and ZsGreen fluorescence was analysed by CytoflexS.

### Statistics

Values are expressed as mean ± standard deviation as indicated. All statistical analysis was carried out in Prism 8.0 (GraphPad), using one-way ANOVA. P-values below 0.05 were considered significant. In multi-group comparisons, multiple testing correction for pairwise tests among groups was applied using Tukey’s post hoc analysis.

## Supplementary Information

**Supplementary Figure 1:**
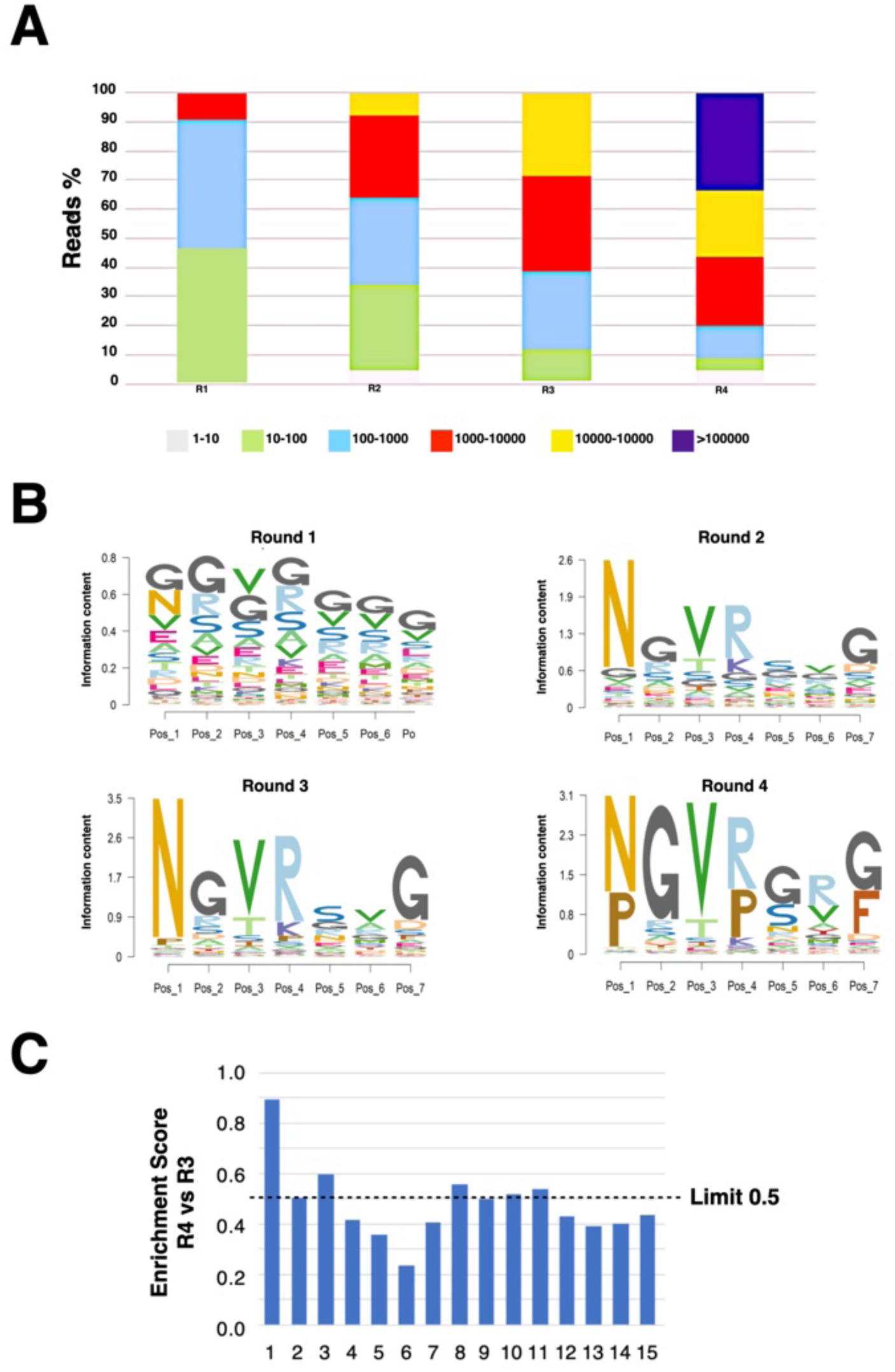
Sequencing analysis of the viral DNA samples isolated from the brain tissue of treated mice. **A)** Clonal distribution of AAV9 library particles subjected to NGS. The relative number of NGS reads was plotted, based on abundance, in six different decimal ranges. **B)** AA frequency logos where each position represented the heptapeptides most frequent residues in the four rounds of selection, in which the library was screened. **C)** Enrichment score as calculated in Korbelin et al. (17) for the target tissue relative to the top fifteen heptapetide that displayed the higher frequency in the fourth round with respect to the third round. The dash line indicates a value of 0.5 which represents heptapeptides presenting the same relative abundance in both rounds. Top 1 and top 2 heptapeptides respectively correspond to AAV-Se2 and AAV-Se1.

**Supplementary Figure 2:**
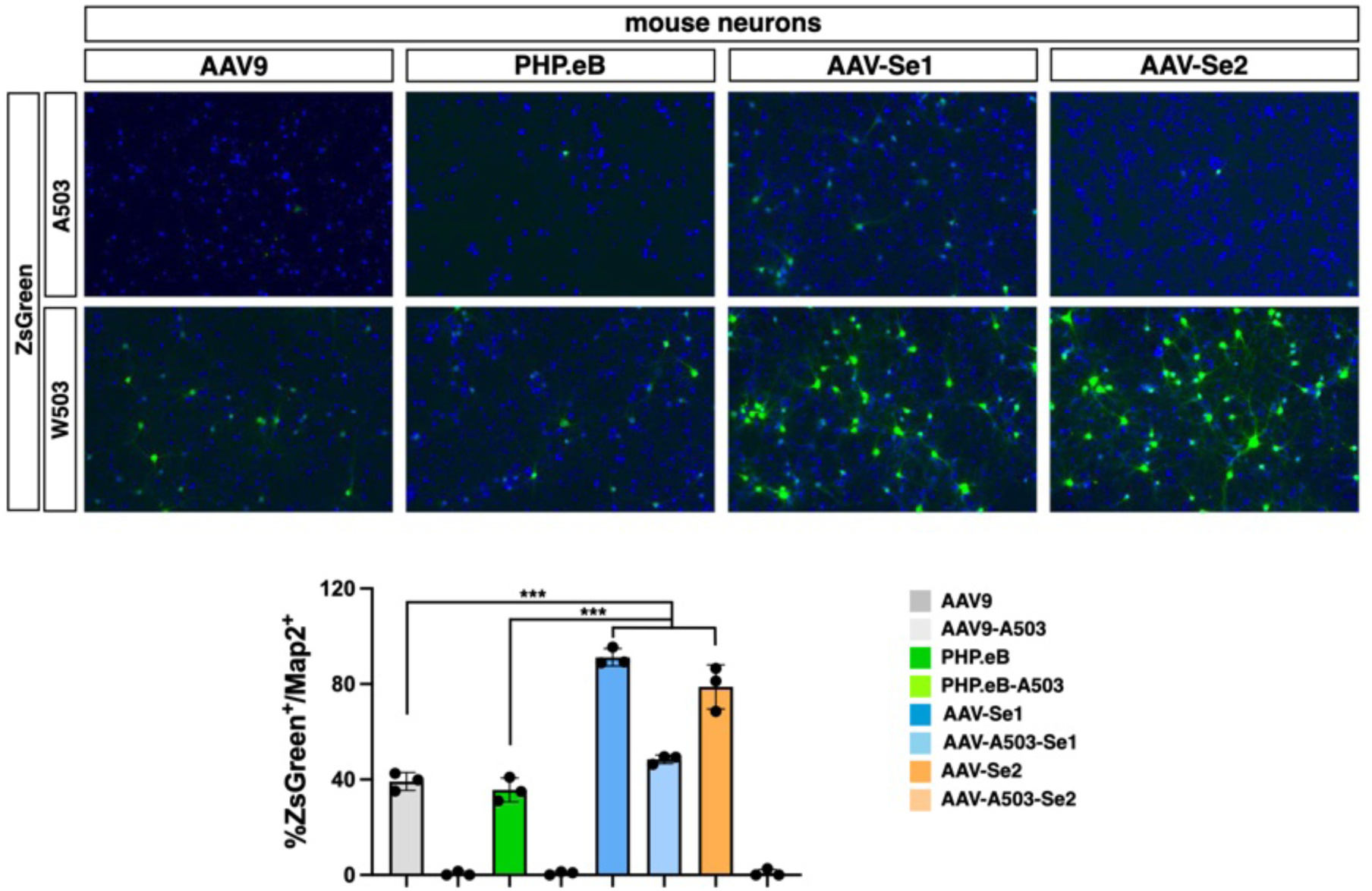
Gene transfer efficiency of viral capsids on mouse primary neuronal cultures. ZsGreen immunofluorescence and its quantification on Map2+ neurons of neuronal cultures infected with AAV9, PHP.eB, AAV-Se1 and AAV-Se2 and their respective A503 mutant variants. Values are mean ± SD. ***p < 0.001. Statistical analysis is performed using one-way ANOVA followed by Tukey post-test.

**Supplementary Figure 3:**
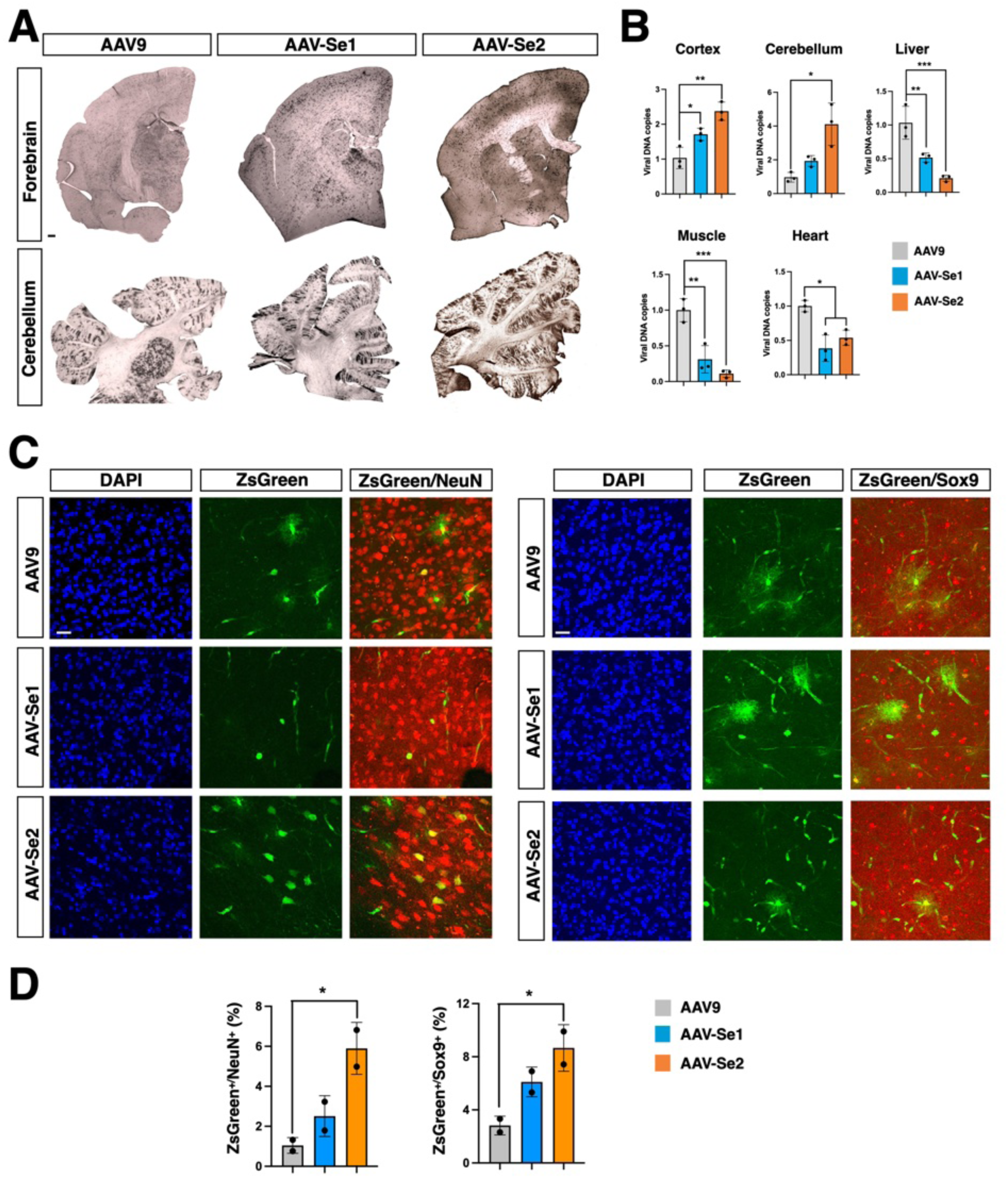
Brain distribution and cell-type specific tropism of AAV9 and AAV-Se1/2 in adult marmosets. **A)** Low magnification images of immunohistochmistry for ZsGreen on forebrain (upper) and cerebellar (lower) sections derived from marmosets transduced with AAV9, AAV-Se1 and AAV-Se2. Scale bars, 200 µm. **B)** Viral DNA vector biodistribution by qRT-PCR in cortex, cerebellum, liver, muscle, heart derived from marmosets treated with AAV9, AAV-Se1 and AAV-Se2. The results were reported as the fold change of viral ZsGreen DNA in marmosets treated with AAV-Se1 and AAV-Se2 relative to marmoset treated with AAV9 (n = 3 pieces for each tissue). **C)** Representative high magnification images of double immunofluorescence stainings for ZsGreen with either the neuronal NeuN or the astrocyte Sox9 markers performed on cerebral cortical sections of marmosets treated with AAV9, AAV-Se1 and AAV-Se2. Scale bars, 100 µm. **D)** Quantitative assessment of neurons (NeuN^+^) and astrocytes (Sox9^+^) in the cortices expressing ZsGreen delivered by AAV9, AAV-Se1 and AAV-Se2 (n=2 different fields of each area). Values are mean ± SD. *p < 0.05; **p < 0.01, ***p < 0.001. Statistical analysis is performed using one-way ANOVA followed by Tukey post-test.

**Supplementary Figure 4:**
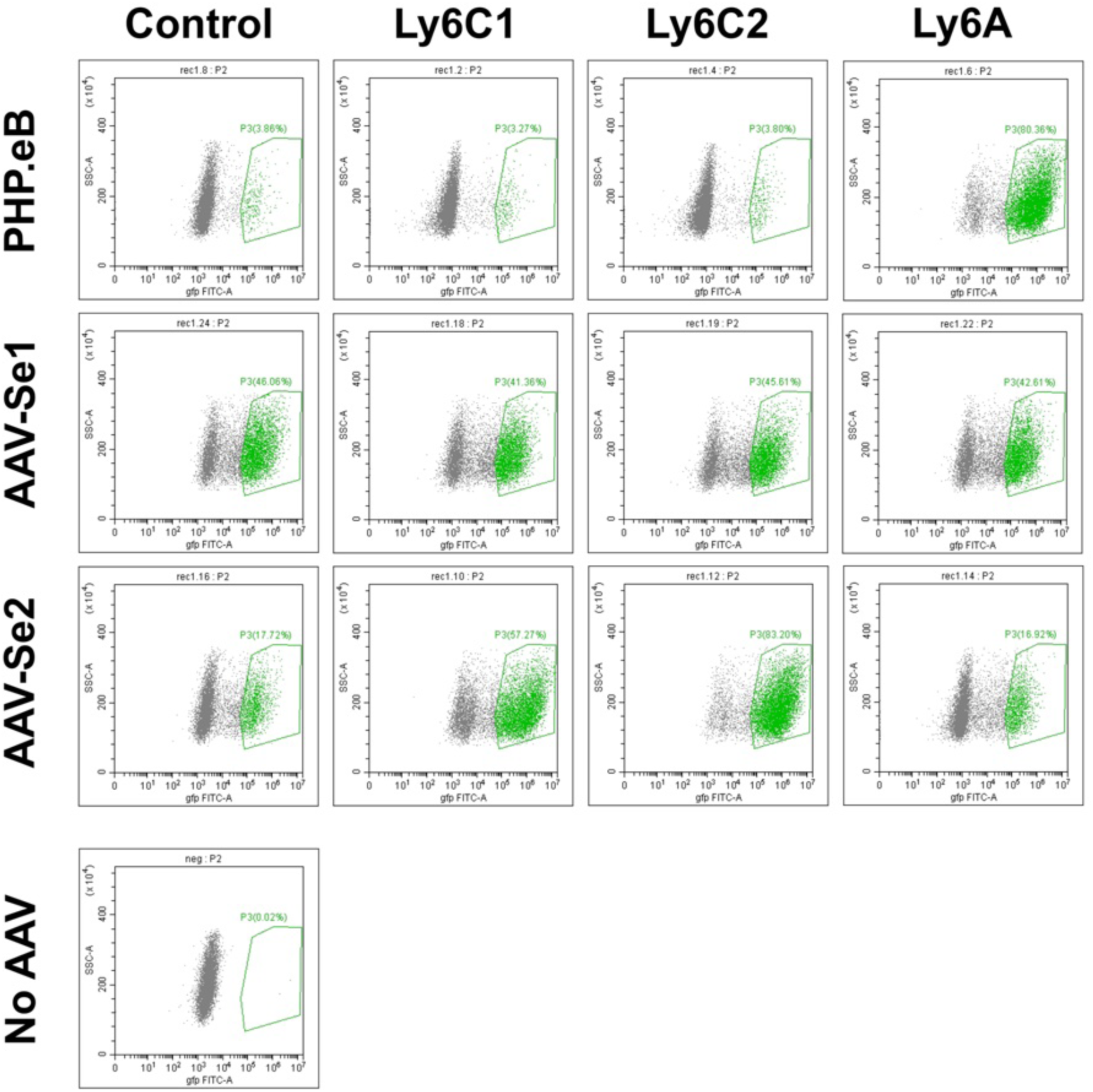
Flow-cytometry plots from transduced HeLa cells. Representative scatter plots of HeLa cells transfected with LY6A, LY6C1, LY6C 2 or untrasfected (control) and subsequently transduced with ZsGreen expressing PHP.eB, AAV-Se1 or AAV-Se2 viral vectors.

**Supplementary Figure 5:**
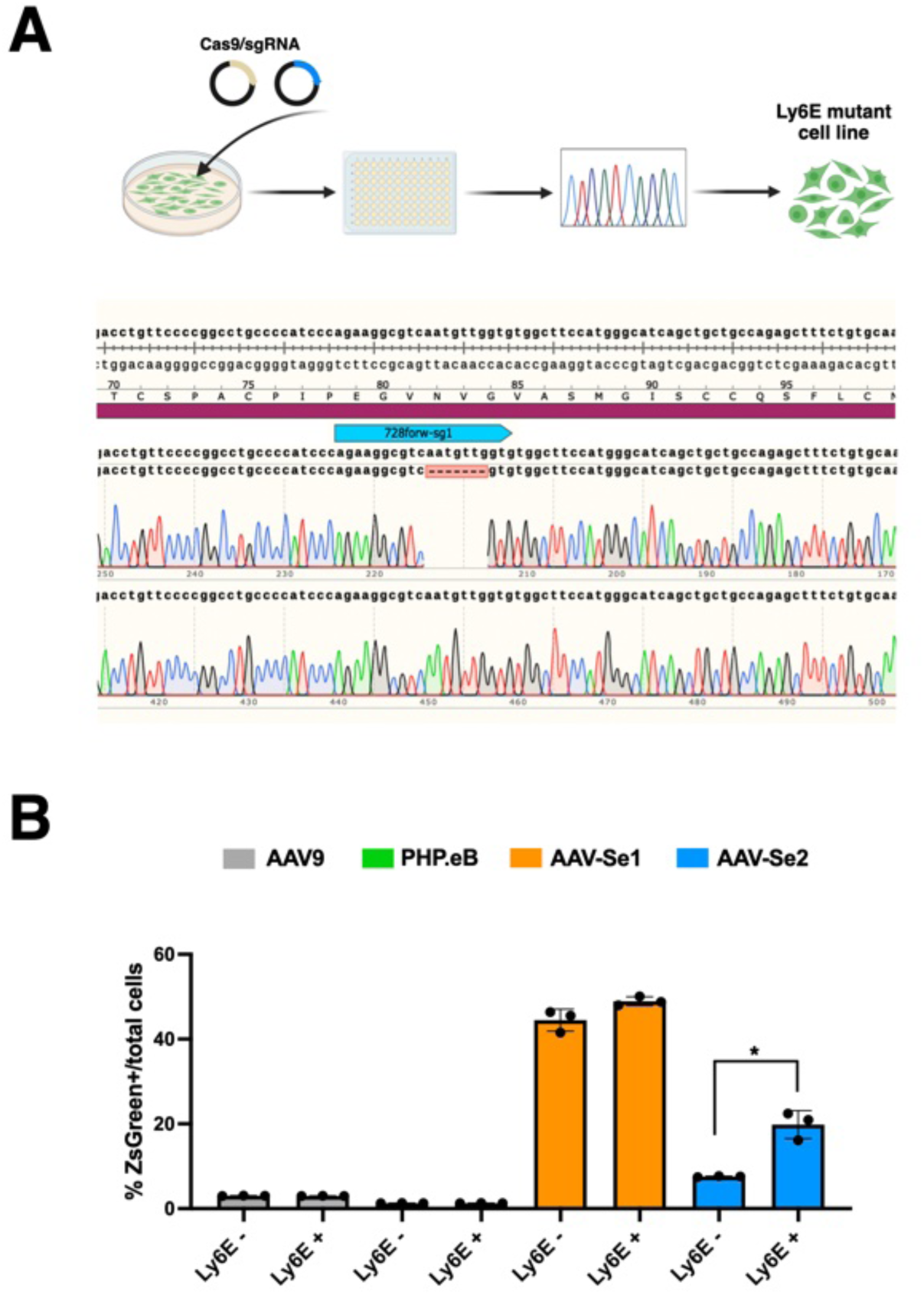
Partial dependence of AAV-Se2 infectivity by LY6E. **A)** Top: Schematic workflow for obtaining HeLa cell clones with targeted inactivation of LY6E by CRISPR/Cas9 gene editing. Bottom: Sanger sequencing results showing a homozygous 7 base deletion in the LY6E coding sequence leading to frameshift and loss of gene activity. **B) D,E**) ZsGreen immunostaining quantification of control or LY6E mutant HeLa cells transduced with ZsGreen expressing AAV9, PHP.eB, AAV-Se1 or AAV-Se2 viral vectors. Only AAV-Se2 transduction is impaired upon loss of LY6E. Values are mean ± SD of n = 3 independent experiments. *p < 0.05; **p < 0.01, ***p < 0.001. Statistical analysis is performed using unpaired *t*-test.

**Flow-cytometry plots from transduced HeLa cells**

Representative scatter plots of HeLa cells transfected with Ly6a, Ly6c1, Ly6c2 expressing plasmids or untrasfected (control) and subsequently transduced with ZsGreen expressing PHP.eB, AAV-Se1 or AAV-Se2 viral vectors.

**Supplementary Table 1: Analysis of basal immunity to AAV capsids of untreated marmosets**

**Supplementary Table 2: Biochemical and cellular analyses of marmoset blood samples**

